# Nanobody generation and structural characterization of *Plasmodium falciparum* 6-cysteine protein Pf12p

**DOI:** 10.1101/2020.12.12.422521

**Authors:** Melanie H. Dietrich, Li-Jin Chan, Amy Adair, Sravya Keremane, Phillip Pymm, Alvin W. Lo, Yi-Chun Cao, Wai-Hong Tham

## Abstract

Surface-associated proteins play critical roles in the *Plasmodium* parasite life cycle and are major targets for vaccine development. The 6-cysteine (6-cys) protein family is expressed in a stage-specific manner throughout *Plasmodium falciparum* life cycle and characterized by the presence of 6-cys domains, which are β-sandwich domains with conserved sets of disulfide bonds. Although several 6-cys family members have been implicated to play a role in sexual stages, mosquito transmission, evasion of the host immune response and host cell invasion, the precise function of many family members is still unknown and structural information is only available for four 6-cys proteins. Here, we present to the best of our knowledge, the first crystal structure of the 6-cys protein Pf12p determined at 2.8 Å resolution. The monomeric molecule folds into two domains, D1 and D2, both of which adopt the canonical 6-cys domain fold. Although the structural fold is similar to that of Pf12, its paralog in *P. falciparum*, we show that Pf12p does not complex with Pf41, which is a known interaction partner of Pf12. We generated ten distinct Pf12p-specific nanobodies which map into two separate epitope groups; one group which binds within the D2 domain, while several members of the second group bind at the interface of the D1 and D2 domain of Pf12p. Characterization of the structural features of the 6-cys family and their associated nanobodies provide a framework for generating new tools to study the diverse functions of the 6-cys protein family in the *Plasmodium* life cycle.

## Introduction

*Plasmodium falciparum* is the most lethal of human malaria species and responsible for the majority of malaria related deaths [1]. One of the key protein families in *P. falciparum* is the 6-cysteine (6-cys) protein family with members representing some of the most abundant surface-expressed proteins across all stages of the malaria parasite life cycle [2]. In *P. falciparum*, there are 14 members in the 6-cys protein family and they share a common structural feature, the 6-cys domain or otherwise referred to as the s48/45 domain. In general, the 6-cys proteins interact with specific human or mosquito proteins for entry into host tissues or to evade the host immune response to promote survival of the malaria parasites [3-6]. Furthermore, several members of the 6-cys proteins are involved in parasite sexual development and fertilization of gametes [7, 8].

The 6-cys proteins are expressed during multiple stages of the parasite life cycle; Pfs230, Pfs48/45, Pfs230p, Pfs47 and PfSOP12 in the sexual stages; Pf52, Pf36, PfLISP2 and PfB9 in the liver stages; and Pf12, Pf12p, Pf41, Pf38, and Pf92 in the blood stages [2]. Pfs230 and Pfs48/45 are the leading transmission blocking vaccine candidates against malaria [9, 10]. Pfs230 forms a complex with Pfs48/45 on the surface of gametocytes, and the complex is involved in the fertilization of male and female gametes [8, 11-13]. Monoclonal antibodies (mAbs) against Pfs230 and Pfs48/45 are effective in blocking transmission by inhibiting successful gamete fertilization [14, 15]. Pfs47 is expressed on the surface of female gametocytes, zygotes and ookinetes [16-18]. The natural selection of specific Pfs47 haplotypes is consistent with the adaptation of *P. falciparum* to different *Anopheles* mosquito species. Through its interaction with a specific mosquito midgut receptor protein, Pfs47 is involved in a lock and key model that drives host tropism between parasite and mosquito [6]. Pf52 and Pf36 are present on the surface of sporozoites and are crucial for invasion of hepatocytes and the formation of a parasitophorous vacuole that envelopes the growing parasite. Entry into liver cells is proposed to involve Pf36 interaction with hepatocyte receptors EphA2, CD81, and Scavenger Receptor BI, which is a critical step for successful malaria infection in the human host [3, 19, 20]. Pf12 and Pf41 form a complex on the merozoite surface and are targets of naturally acquired immunity [21-26]. Pf12 is the fifth most prevalent glycosylphosphatidylinositol (GPI)-anchored protein on the merozoite surface and Pf12 and Pf41 have been implicated to be involved in red blood cell invasion [4, 27]. Pf92 is an abundant merozoite surface protein and constitutes about 5% of the total surface coat [27]. Pf92 plays a role in immune evasion by recruiting human complement regulator Factor H, which is the major complement regulator of the alternative pathway of complement. This recruitment serves to downregulate complement activation on the merozoite surface and protect *P. falciparum* merozoites from complement-mediated lysis [5].

The 6-cys domain has two to six cysteines that form disulfide bonds and is evolutionary related to the SAG1-related sequence (SRS) domain in *Toxoplasma gondii* [28]. The 6-cys and SRS domains have been proposed to be derived from an ephrin-like precursor originating from a vertebrate host protein, with a general function to mediate extracellular protein-protein interactions and cellular adhesion [28]. The 6-cys domain is characterized by a β-sandwich fold of parallel and anti-parallel β-strands and a conserved cysteine motif [28-30]. The β-sandwich is formed by two β-sheets, usually termed A and B, that are pinned together by two disulfide bonds. A third disulfide bond connects a loop region to the core structure and a small β-sheet of two β-strands runs perpendicular along the side of β-sheet B [24, 26, 28-30]. Between one to fourteen 6-cys domains (denoted as D1, D2, D3…) are present in each 6-cys protein and they are often found in tandem pairs of A- and B-type 6-cys domains [26, 30]. The two types of 6-cys domains differ in the number of β-strands in β-sheet A, with A-type domains usually containing four and B-type domains usually containing five β-strands. The position and connectivity of cysteines differ between the 6-cys domain compared to the SRS-domain in *T. gondii*. In the 6-cys domain the disulfide bond connectivity follows a C1-C2, C3-C6, C4-C5 pattern, whereas in the SRS domain it follows a C1-C6, C2-C5, and C3-C4 pattern [31-33]. Domains containing less than six cysteines have been identified in both protein families [34-37].

Four crystal structures have been determined for the 6-cys family members; the C-terminal D3 domain of Pfs48/45 bound to either inhibitory mAb 85RF45.1 or mAb TB31F (humanized version of 85RF45.1) [38, 39], the D1 and D2 domains of Pf41 [24], the D1 and D2 domains of Pf12 [26], and the D1 domain of Pfs230 with transmission-blocking mAb 4F12 [40]. While these structures have been important in elucidating the general fold of the 6-cys domain and inhibitory epitopes for two transmission-blocking antibodies, high-resolution crystal structures of all 6-cys proteins will be required to fully understand the diverse functions of this protein family. Alpacas, llamas and their camel cousins have evolved one of the smallest naturally occurring antigen recognition domains called nanobodies. Nanobodies are ∼15 kDa in size, display strong binding affinities to target proteins and also function as structural chaperones to assist in crystal formation. Nanobodies may be used both to assist in the crystallisation of 6-cys proteins and to block malaria parasite invasion by inhibiting the specific functions of 6-cys proteins.

While several family members play critical roles in the parasite life cycle, many 6-cys proteins are not well characterized and their precise functions are unknown. One of the understudied 6-cys protein is Pf12p, which is a paralog of Pf12 [30]. Pf12p is predicted to contain a signal peptide, two 6-cys domains and a GPI-anchor that links the protein to the parasite surface [27]. Microarray data indicates that Pf12p is transcribed in blood stages and mass spectrometry data suggests that this protein is also present in sporozoites but the function of Pf12p is currently unknown [27, 41, 42]. To further characterize Pf12p using structural methods and to generate antibody tools that are specific to Pf12p, we immunized an alpaca with recombinant Pf12p protein to produce specific nanobodies. We characterized a collection of anti-Pf12p nanobodies for their specificity, affinities and epitope competition. To the best of our knowledge, we determined the first high-resolution crystal structures of Pf12p alone and Pf12p bound to two distinct nanobodies. These crystal structures and nanobody tools will help to drive functional analyses of Pf12p in the future.

## Materials and Methods

### Expression and purification of Pf12p, Pf12 and Pf41

We expressed recombinant fragments of Pf12p corresponding to amino acids N24-S341 (Pf12p D1D2) and N168-S341 (Pf12p D2). The baculovirus transfer vector pAcGP67-A was modified to introduce a Tobacco etch virus (TEV)-cleavable His_8_-tag following the GP-67 signal sequence. The Pf12p sequences were cloned after the N-terminal TEV-cleavage site, using NheI and NotI restriction sites. Pf12p proteins were expressed using *Spodoptera frugiperda* (Sf) 21 cells (Life Technologies) cultured in Insect-XPRESS Protein-free Insect Cell Medium supplemented with L-glutamine (Lonza). A Sf21 cell culture of ∼1.8 × 10^6^ cells/ml was inoculated with the third passage stock of virus and incubated for three days at 28 °C. Cells were separated from the supernatant by centrifugation. The supernatant was concentrated via tangential flow filtration using a 10 kDa molecular weight cut-off cassette (Millipore). The concentrated supernatant was dialyzed into 30 mM Tris pH 7.5, 300 mM NaCl (buffer A) and incubated with Ni-NTA resin (Qiagen) for one hr at 4 °C on a roller shaker. The Ni-NTA resin was added onto a gravity flow chromatography column and washed with 10-20 column volumes of buffer A. The imidazole concentration in buffer A was increased stepwise from 0-300 mM for protein elution. TEV protease was added to the pooled fractions containing Pf12p and dialysed into buffer A. The solution was incubated with Ni-NTA resin (Qiagen) for one hr at 4 °C to bind the His-tagged TEV protease and uncleaved Pf12p. Untagged Pf12p was collected from the flowthrough, concentrated and applied onto a size exclusion chromatography (SEC) column (SD200 increase 10/300 or SD200 16/600 pg, GE Healthcare) pre-equilibrated with 20 mM HEPES pH 7.5, 150 mM NaCl.

Recombinant fragments of Pf12 D1D2 (residues N28-S304) and Pf41 D1D2 (residues K21-S368) were cloned into our modified pAcGP67-A vector, expressed and purified following the same purification protocol as Pf12p above with some modifications.

### Alpaca Immunisation and nanobody phage library

One alpaca was subcutaneously immunized six times 14 days apart with approximately 200 μg of recombinant Pf12p D1D2 protein. The adjuvant used was GERBU FAMA. Immunization and handling of the alpaca for scientific purposes was approved by Agriculture Victoria, Wildlife & Small Institutions Animal Ethics Committee, project approval No. 26-17. Blood was collected three days after the last immunization for the preparation of lymphocytes. Nanobody library construction was carried out according to established methods [43]. Briefly, alpaca lymphocyte mRNA was extracted and amplified by RT-PCR with specific primers to generate a cDNA library size of 10^8^ nanobodies with 80% correct sized nanobody insert. The library was cloned into a pMES4 phagemid vector amplified in *Escherichia coli* TG1 strain and subsequently infected with M13K07 helper phage for recombinant phage expression.

### Isolation of Pf12p nanobodies

Biopanning for Pf12p nanobodies using phage display was performed as previously described [43]. Phages displaying Pf12p-specific nanobodies were enriched after two rounds of biopanning on 1 μg of immobilized Pf12p D1D2 protein. After the second round of panning, 95 individual clones were selected for further analyses by ELISA for the presence of Pf12p nanobodies. Positive clones were sequenced and annotated using the International ImMunoGeneTics database (IMGT) and aligned in Geneious Prime.

### Expression and purification of nanobodies

Nanobodies were expressed in *E. coli* WK6 cells. Bacteria were grown in Terrific Broth at 37 °C to an OD_600_ of 0.7, induced with 1 mM IPTG and grown overnight at 28 °C for 16 h. Cell pellets were harvested and resuspended in 20% sucrose, 20 mM imidazole, 150 mM NaCl DPBS and incubated for 15 min on ice. 5 mM EDTA was added and incubated on ice for 20 minutes. After this incubation, 10 mM MgCl_2_ was added and periplasmic extracts were harvested by centrifugation and the supernatant was loaded onto a 1 ml HisTrap FF column (GE Healthcare). The nanobody was eluted via a linear gradient into 400 mM imidazole, 100 mM NaCl, PBS. The appropriate fractions were concentrated and subjected to SEC (SD200 increase 10/300) pre-equilibrated in 20 mM HEPES pH 7.5, 150 mM NaCl.

### ELISA for antibody specificity

96-well flat-bottomed MaxiSorp plates were coated with 65 nM of recombinant protein as indicated in 50 μL of PBS at room temperature (RT) for one hour. All washes were done three times using PBS and 0.1% Tween (DPBS-T) and all incubations were performed for one hour at RT. Coated plates were washed and blocked by incubation with 10% skim milk solution. Plates were washed and then incubated with 65 nM of nanobodies. The plates were washed and incubated with mouse anti-His (Bio-Rad MCA-1396; 1:1000) followed by horseradish peroxidase (HRP)-conjugated goat anti-mouse secondary antibody (MerckMillipore AP124P, 1:1000). After a final wash, 50 μL of azino-bis-3-ethylbenthiazoline-6-sulfonic acid (ABTS liquid substrate; Sigma) was added and incubated in the dark at RT and 50 μL of 1% SDS was used to stop the reaction. Absorbance was read at 405 nm and all samples were done in duplicate.

### Western Blotting

Purified Pf12p D1D2 was loaded on a 4-12% Bis-Tris SDS-PAGE gel under reduced and non-reduced conditions, and proteins were transferred onto a PVDF membrane. All washes were done in DPBS-T at RT for five minutes. The membrane was blocked with 10% milk in DPBS-T overnight at 4 °C. The membrane was washed and incubated with 0.5 ug/ml nanobody in 1% milk in DPBS-T for one hour at RT. The membrane was washed twice and incubated with HRP-conjugated goat anti-llama IgG (Agrisera AS10 1240, 1:2000) in DPBS-T for one hour at RT. The membrane was washed twice followed by a final wash with DPBS for 10 min. The blots were processed with an enhanced chemiluminescence (ECL) system (Amersham Biosciences).

### Bio-Layer interferometry (BLI)

Affinity determination measurements were performed on the Octet RED96e (FortéBio). All assays were performed using NiNTA capture sensor tips (NTA) sensors (FortéBio) with kinetics buffer (PBS pH 7.4 supplemented with 0.1% (w/v) BSA and 0.05% (v/v) TWEEN-20) at 25 °C. After a 60 s biosensor baseline step, nanobodies (5 μg/mL) were loaded onto NTA sensors by submerging sensor tips until a response of 0.5 nm and then washed in kinetics buffer for 60 s. Association measurements were performed using a two-fold dilution series of untagged Pf12p D1D2 from 6-200 nM (used a dilution series from 16-500 nM to measure affinity to nanobody A10) for 180 s and dissociation was measured in kinetics buffer for 180 s. Sensor tips were regenerated using a cycle of 5 s in 300 mM imidazole pH 7.5 and 5 s in kinetics buffer repeated five times. Baseline drift was corrected by subtracting the response of a nanobody loaded sensor not incubated with untagged Pf12p D1D2. Curve fitting analysis was performed with Octet Data Analysis 10.0 software using a global fit 1:1 model to determine K_D_ values and kinetic parameters. Curves that could not be fitted were excluded from the analyses. Mean kinetic constants reported are the result of three independent experiments.

The affinity of Pf12 and Pf12p binding to Pf41 were measured using the method above with the following modifications. His-tagged Pf41 (10 μg/mL or 20 μg/mL) was loaded onto NTA sensors until a response shift of 1.8 nm. Association measurements were performed using a two-fold dilution series from 16-500 nM (if loaded with 10 μg/mL of Pf41) or 31-1000 nM (if loaded with 20 μg/mL of Pf41) of untagged Pf12 D1D2 and untagged Pf12p D1D2.

### Competition binding experiment using BLI

For competition experiments using BLI, 150 nM untagged Pf12p D1D2 was pre-incubated with each nanobody at a 10-fold molar excess for one hr at RT. A 30 s baseline step was established between each step of the assay. NTA sensors were first loaded with 10 μg/mL of nanobody for 5 min. The sensor surface was then quenched by dipping into 20 μg/mL of an irrelevant nanobody for 5 min. Nanobody-loaded sensors were then dipped into premixed solutions of Pf12p D1D2 and nanobody for 5 min. Nanobody-loaded sensors were also dipped into Pf12p D1D2 alone to determine the level of Pf12p D1D2 binding to immobilized nanobody in the absence of other nanobodies. Percentage competition was calculated by dividing the max response of the premixed Pf12p D1D2 and nanobody solution binding by the max response of Pf12p binding alone, multiplied by 100.

### Crystallization and Structure Determination

Purified Pf12p was mixed with an anti-Pf12p nanobody in a molar ratio of 1:1.5 and incubated for 1 hr on ice prior to SEC (SD200 increase 10/300; 20 mM HEPES pH 7.5, 150 mM NaCl) to separate the Pf12p-nanobody complex from excess nanobody. Crystallization trials were performed at the Collaborative Crystallization Centre (CSIRO, C3, Parkville) at 8 °C. Hanging drop vapour diffusion crystallization trials were set up in-house for crystal optimization of Pf12p-D9 and Pf12p-B9. Pf12p crystals were obtained in 12.5% MPD, 0.02 M alanine, 0.1 M bicine-tris pH 8.5, 0.02 M glycine, 0.02 M lysine, 12.5% PEG1000, 12.5% PEG3350, 0.02 M serine, 0.02M sodium glutamate, 0.2 M magnesium chloride at 4 mg/ml and harvested in mother liquor. Pf12p-D9 crystallized in 0.1 M bis-tris chloride pH 6.5, 0.2 M magnesium chloride, 25% PEG3350 at 5 mg/ml and were flash frozen in mother liquor containing 30% glycerol. Pf12p-B9 crystallized in 20% PEG3000, 0.1 M trisodium citrate-citric acid pH 5.5 at 6 mg/ml. A Pf12p-B9 crystal was transferred stepwise into cryo-protectant and soaked for 3 min in mother liquor containing 3 mM K_2_Pt(CN)_4_ and 25% glycerol before flash-freezing in liquid nitrogen. X-ray diffraction data was collected at the MX2 beamline at the Australian Synchrotron. The XDS package [44] was used for data processing. Phaser [45] was used for molecular replacement. The phase problem of Pf12p-B9 was solved using coordinates of nanobody VHH-72 of PDB ID 6WAQ, and modified structures of an unpublished 6-cys protein and Pf12 (PDB ID 2YMO) as model structures. The 6-cys protein models were trimmed to remove flexible loops and amino acids other than cysteines were modified to alanine. Initial phases of Pf12p and Pf12p-D9 were obtained using coordinates of individual chains of Pf12p-B9 as search models for molecular replacement. Alternating rounds of structure building and refinement were carried out using *Coot* [46] and phenix [47, 48]. About 2000 reflections were set aside in each case for the calculation of R_free_. Figures of the structures were prepared with PyMOL (www.pymol.com) [49]. Interactions, interfaces and buried areas from solvent were analysed using PISA [50]. The atomic coordinates and structure factor files have been deposited in the Protein Data Bank (PDB) under PDB ID 7KJ7 for Pf12p, 7KJH for Pf12p-B9 and 7KJI for Pf12p-D9.

### Size Exclusion Chromatography Binding Studies of Pf12p with Pf12 and Pf41

Complexation was carried out by incubating 100 μg Pf12p or Pf12 with Pf41 at a 1:1 molar ratio for one hr at RT. 30 μL of the sample was loaded onto an SEC column (Superdex 200 3.2/300) pre-equilibrated in 20 mM HEPES pH 7.5, 150 mM NaCl using a 100 μL loop. The run was carried out using a 0.03 ml/min flow rate and 100 μL fraction size. Equivalent amounts of Pf12, Pf12p and Pf41 were run singly for comparison of retention volumes to assess complex formation.

## Results

### Isolation and characterization of Pf12p-specific nanobodies

A 10^8^ nanobody phage display library was generated from an alpaca immunised with recombinant Pf12p D1D2 and used to select for Pf12p-specific nanobodies. After two rounds of bio-panning, we identified ten distinct nanobody clonal groups based on differences in the amino acid sequence of the complementary determining region 3 (CDR3) (Figure 1A). The CDR3 regions of the nanobodies vary in at least one amino acid with lengths between 8 to 21 residues. One member of each clonal group was selected for further characterization and will be referred to as A10, B2, B9, B12, C4, C12, D9, F7, G6, and H7. These nanobodies were expressed and purified with overall yields of 1-12 mg per litre of initial culture and migrated between 13 and 17 kDa on SDS-PAGE under reducing conditions (Figure 1B).

**Fig 1.**
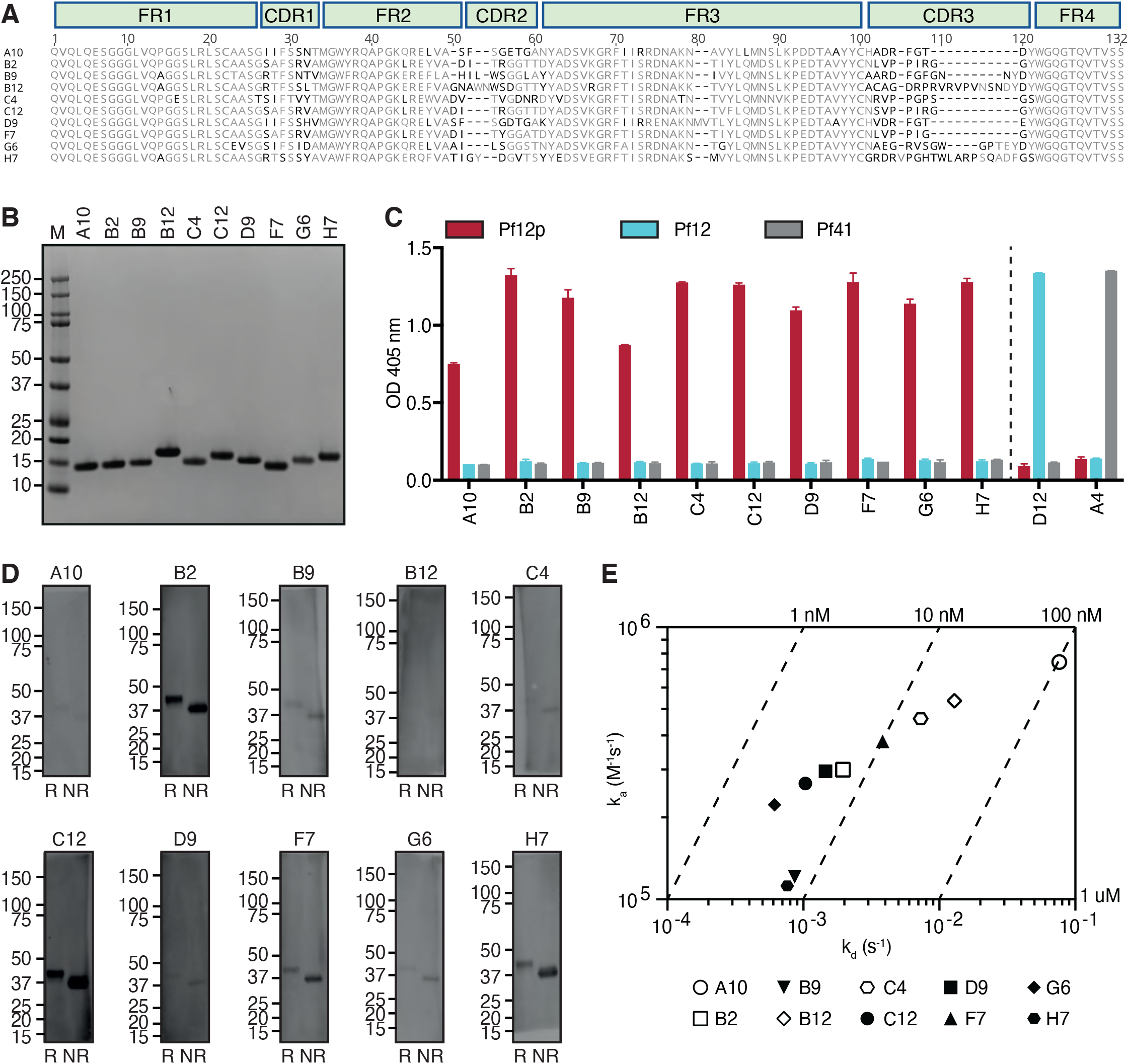
Pf12p-specific nanobodies. **(A)** Sequence alignment of 10 nanobodies with framework regions (FR) and complementary determining regions (CDR) indicated according to the international ImMunoGeneTics information system (IMGT). Black residues represent less than 60% similarity to the consensus sequence. **(B)** Coomassie-stained SDS-PAGE gel of purified Pf12p-specific nanobodies under reducing conditions. Molecular weight marker (M) in kDa is shown on the left-hand side. **(C)** Detection of Pf12p by nanobodies using ELISA. Anti-Pf12p nanobodies, anti-Pf12 nanobody D12 and anti-Pf41 nanobody A4 were added to microtiter wells coated with Pf12p, Pf12 and Pf41. Bound nanobodies were detected with anti-His antibody followed by HRP-conjugated secondary antibody. Error bars represent standard deviation of the mean. **(D)** Detection of Pf12p by nanobodies by Western blotting. Reduced (R) and non-reduced (NR) Pf12p protein was separated by SDS-PAGE and probed with the respective nanobodies and detected using an HRP-conjugated goat anti-llama IgG. Molecular weight marker in kDa is shown on the left hand-side. **(E)** Iso-affinity plot showing the dissociation rate constants (k_d_) and association rate constants (k_a_) of Pf12p nanobodies as measured by BLI. Symbols that fall on the same diagonal lines have the same equilibrium dissociation rate constants (K_D_) indicated on the top and right sides of the plot.

To examine the specificty of these Pf12p-specific nanobodies, we used recombinant Pf12p and two other recombinant 6-cys proteins, Pf12 and Pf41 in an ELISA-based assay (Figure 1C). All three recombinant proteins consist of two 6-cys domains. Pf12p shares 16.7% sequence identity with Pf12 and 12.8% with Pf41. All ten nanobodies recognize Pf12p but do not bind to Pf12 nor Pf41. Pf12- and Pf41-specific nanobodies, D12 and A4, respectively, do not cross-react with Pf12p. Collectively, these results show that the ten nanobodies are specific to Pf12p and are not cross-reactive with two other 6-cys proteins.

We wanted to determine whether the Pf12p-specific nanobodies are able to detect Pf12p by Western blotting under reducing and non-reducing conditions (Figure 1D). Six of the ten nanobodies, A10, B9, B12, C4, D9, and G6 showed no or weak reactivity under both conditions. Four nanobodies, B2, C12, F7, and H7, recognize the reduced and non-reduced Pf12p to different extents. All four nanobodies above show a stronger signal with non-reduced Pf12p compared to reduced protein, indicating that the presence of disulfide bonds improves the recognition of Pf12p by these nanobodies using Western blotting.

We used bio-layer interferometry (BLI) to determine the binding kinetics and affinities of the interaction between nanobodies and Pf12p (Figure 1E and Supplementary Figure S1). Nine out of ten nanobodies bind recombinant Pf12p with high affinity in the low nanomolar range, with association rates around 10^5^ M^-1^s^-1^ and dissociation rates between 10^−2^ and 10^−4^ s^-1^. A10 which is the weakest binding nanobody has an affinity of ∼100 nM.

### Pf12p-specific nanobodies bind to two separate regions on Pf12p

To determine whether the Pf12p-specific nanobodies bound epitopes within the D1 or D2 domain of Pf12p, we performed an ELISA using recombinant Pf12p D1D2 and Pf12p D2 proteins. Our recombinant Pf12p D1D2 protein contains both predicted 6-cys domains and lacks the N-terminal signal sequence and predicted C-terminal GPI-anchor (Figure 2A). Pf12p D2 contains the C-terminal domain D2 only (Figure 2A). Unfortunately, we were unable to express the single domain D1 of Pf12p. Nanobodies B2, C4, C12, F7, and H7, bound to both Pf12p D1D2 and Pf12p D2 proteins with similar binding signals showing that these five nanobodies bind epitopes within the D2 domain of Pf12p (Figure 2B). Nanobodies B9 and G6 showed a lower detection signal to Pf12p D2 compared to Pf12p D1D2, suggesting that both the D1 and D2 domains of Pf12p may be involved in nanobody binding. A10, B12, and D9, bound Pf12p D1D2 but their signal for Pf12p D2 was weaker or similar to that of the negative controls (Figure 2B). The binding sites of these three nanobodies may lay within the D1 domain, but a contribution of D2 cannot be excluded.

**Fig 2.**
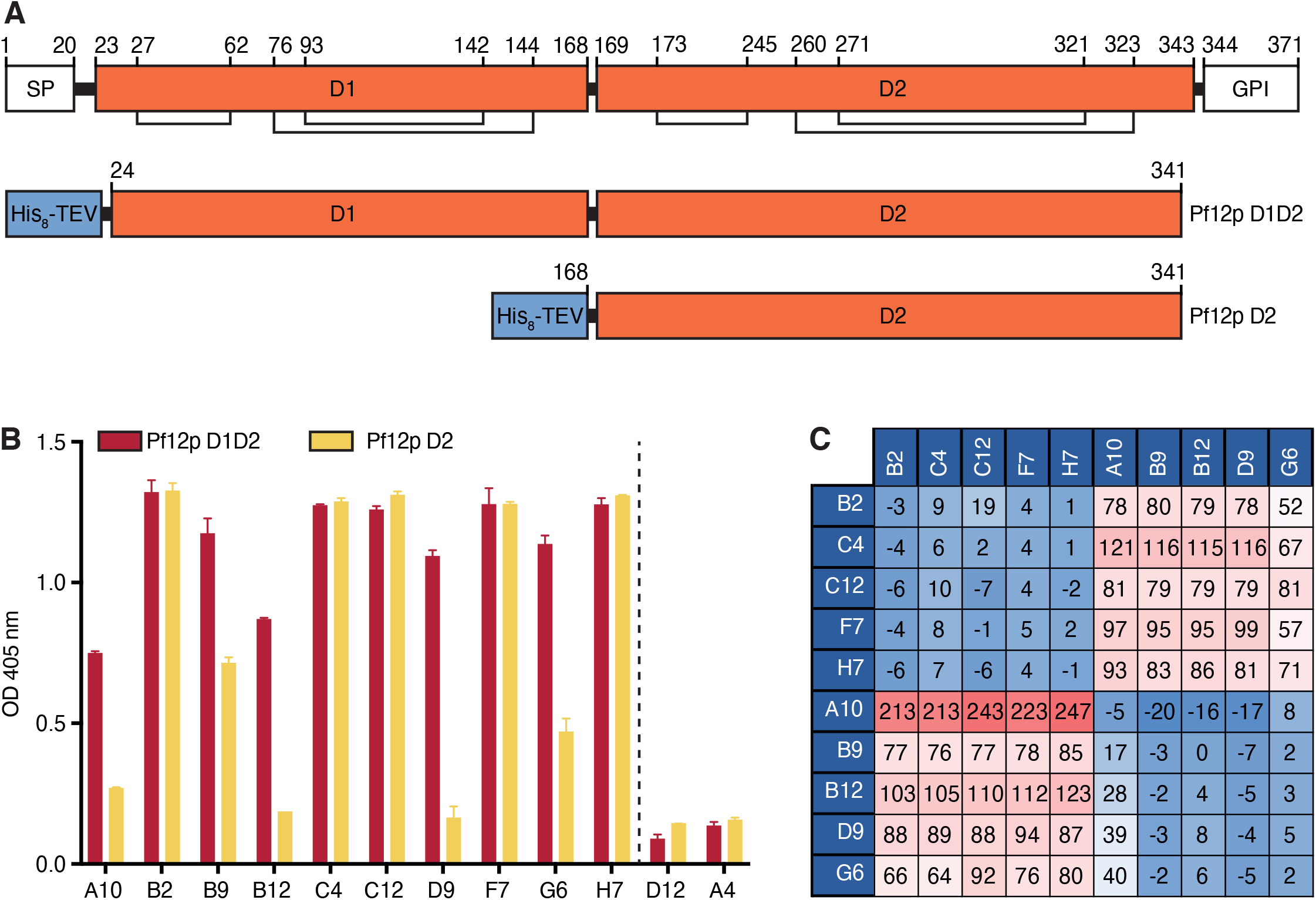
Domain mapping and epitope competition of Pf12p-specific nanobodies. **(A)** Domain organization of full-length Pf12p (upper) and recombinant fragments of Pf12p D1D2 (middle) and Pf12p D2 (lower). SP, signal peptide; GPI, predicted GPI-anchor sequence; His_8_-TEV, N-terminal His_8_-tag followed by a TEV cleavage site. Lines and numbers show cysteine bonds. **(B)** Domain mapping of Pf12p-specific nanobodies using ELISA. Anti-Pf12p nanobodies were added to microtiter wells coated with Pf12p D1D2 and Pf12p D2. Bound nanobodies were detected with anti-His antibody followed by HRP-conjugated secondary antibody. Error bars represent standard deviation of the mean. **(C)** Epitope competition experiments by BLI using immobilized nanobodies indicated on the left column incubated with nanobodies indicated on the top row pre-incubated with Pf12p using a 10:1 molar ratio. Binding of Pf12p premixed with nanobody was calculated relative to Pf12p binding alone, which was assigned to 100%. A blue to red gradient shows antibodies with the highest levels of competition in blue and the lowest in red.

To determine if the Pf12p-specific nanobodies recognize similar epitopes, we performed a nanobody competition experiment using BLI. As expected, all nanobodies were able to compete with themselves (Figure 2C). We observed that B2, C4, C12, F7, and H7 compete with each other, while A10, B9, B12, D9 or G6 do not compete with the former set of nanobodies. Consistent with these results, A10, B9, B12, D9 and G6 compete with each other whereas B2, C4, C12, F7 and H7 do not compete with them. Our BLI results show that the nanobodies group into two different epitope bins. Together with our ELISA analysis, we propose that one group B2, C4, C12, F7, and H7 bind within the D2 domain of Pf12p, whereas the binding sites of the other group of competing nanobodies, A10, B9, B12, D9, and G6, involve the D1 domain to differing extents.

### The crystal structure of Pf12p

We determined the crystal structure of Pf12p D1D2 at a resolution of 2.8 Å by molecular replacement. Two molecules are present in the asymmetric unit of our crystal structure, which are nearly identical and align with a root mean square deviation (RMSD) of 0.4 Å. Evaluation of the interfaces using PISA [50] indicates that the protein is monomeric, which is consistent with the elution profile from size exclusion chromatography (Figure 4C). In the following structural description, we will focus solely on molecule A. Our crystal structure reveals that Pf12p D1D2 folds into two domains, D1 and D2, each containing six cysteines (Figure 3A). The N-terminal D1 domain adopts the fold of a typical 6-cys domain of type A, which forms a β-sandwich with a 4-on-4 β-strand arrangement. The two sheets of the β-sandwich consist of mixed parallel and anti-parallel β-strands and are pinned together by two disulfide bonds, formed between C27 and C62, and between C76 and C144. A third disulfide bond between residues C93 and C142 connects a loop to the core structure. In our structure, the cysteines form C1-C2, C3-C6 and C4-C5 pairings, which is characteristic for a typical 6-cys domain.

**Fig 3.**
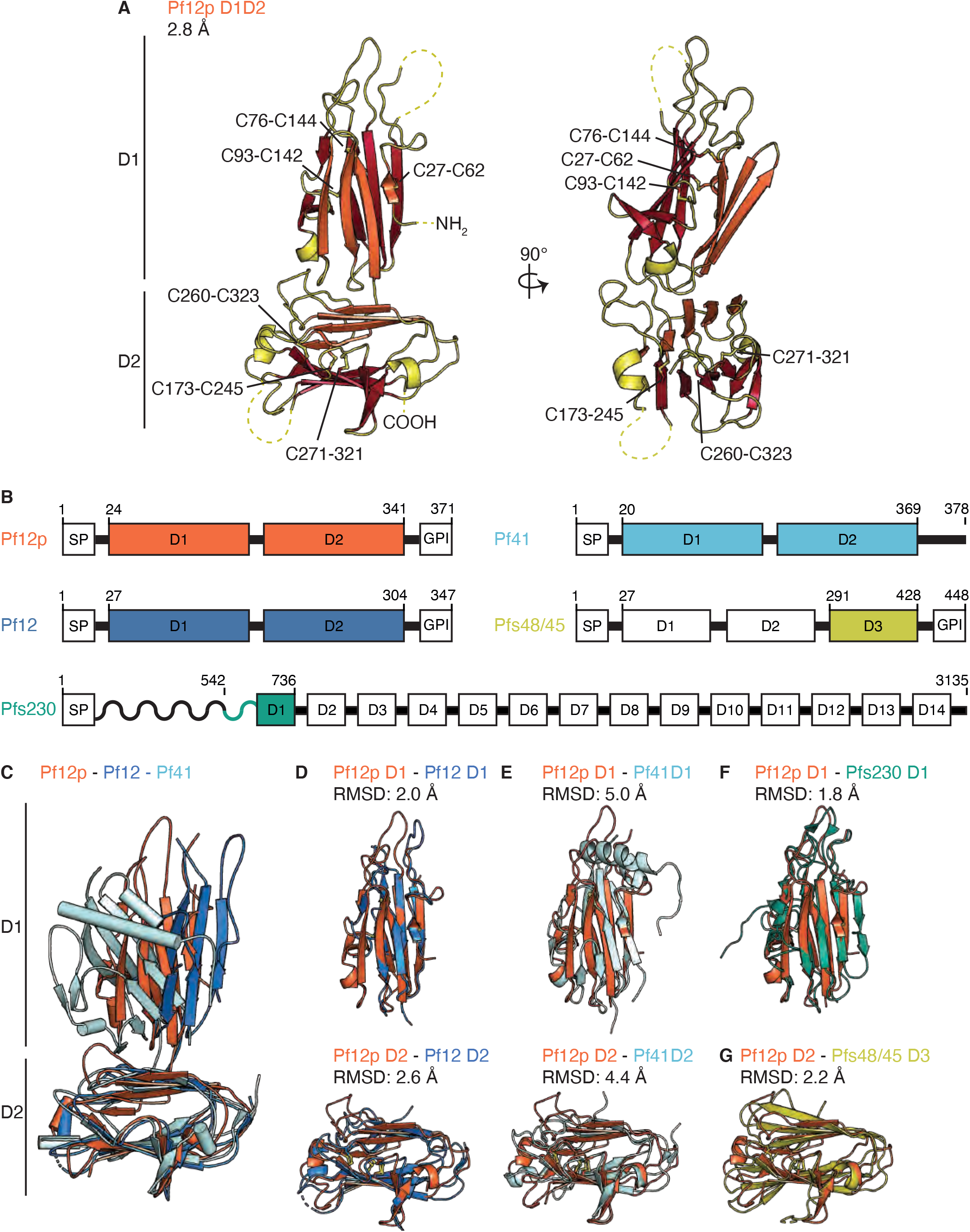
Crystal structure of Pf12p and comparison with structures of other 6-cys protein family members. (**A)** The Pf12p structure (PDB ID 7KJ7) is shown in two orthogonal views. The N- and C-termini and disulfide bonds are labelled. Dashed lines indicate regions which do not have defined electron density. The β-sheets A and B of the β-sandwich of each domain are coloured in orange and red, respectively. **(B)** Schematic diagram of selected 6-cys proteins (not to scale). Predicted 6-cys domains are in white and labelled sequentially. The recombinant fragments used in published structural studies are coloured. SP, signal peptide; GPI, GPI-anchor. Residue numbers are indicated on top. **(C)** Structural alignment of Pf12 (PDB ID 2YMO) and Pf41 (PDB ID 4YS4) with Pf12p based on the D2 domain. **(D)** Superimposition of Pf12p and Pf12 D1 domains (upper) and the D2 domains (lower). **(E)** Superimposition of Pf12p and Pf41 D1 domains (upper) and the D2 domains (lower). **(F)** Superimposition of the D1 domains of Pf12p and Pfs230 (D1M construct) (PDB ID 6OHG). **(G)** Superimposition of the D2 domain of Pf12p with the D3 domain (6C construct) of Pfs48/45 (PDB ID 6E63).

**Fig 4.**
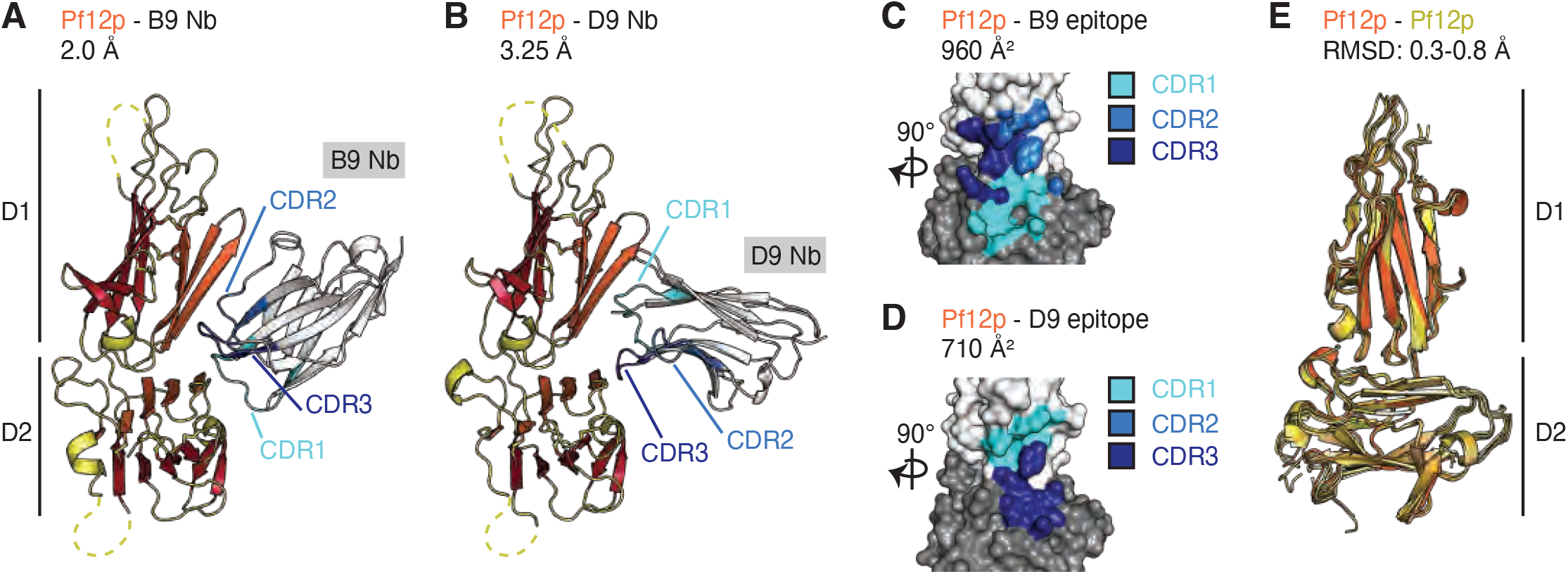
Pf12p does not interact with Pf41. (**A)** BLI-binding experiment with immobilized Pf41 and Pf12 in solution. Representative binding curves of five different Pf12 concentrations are plotted and fitted to a 1:1 binding model. (**B)** BLI-binding experiment with immobilized Pf41 and Pf12p in solution. Six different Pf12p concentrations ranging from 16 - 500 nM were tested, but no binding could be detected. **(C)** SEC analyses show that recombinant Pf12 and Pf41 form a heterodimer. The Pf12-Pf41 complex elutes at a retention volume corresponding to higher molecular weight compared to the individual proteins on SEC. Excess of Pf12 elutes as a second peak at the expected retention volume. **(D)** SEC analyses show that recombinant Pf12p does not form a stable complex with Pf41. The mix of protein elutes at a retention volume between the peak maxima of the individual proteins. Retention volume of molecular weight marker proteins and their corresponding size are indicated.

The C-terminal D2 domain of Pf12p folds into a 6-cys domain of type B forming a β- sandwich with a 5-on-4 β-strand arrangement of mixed parallel and anti-parallel β-strands (Figure 3A). As in D1, three disulfide bonds are present in this domain with C1-C2, C3-C6, and C4-C5 pairs formed by C173-245, C260-323, and C173-C245. While most residues are well resolved, one loop in each domain is partly disordered, namely residues 40-56 in D1 and residues 199-241 in D2, which features an asparagine-rich region which is not conserved in Pf12p homologs in other *Plasmodium* species (Supplementary Figure S2).

The highest structural similarity to Pf12p D1D2 are structures of the merozoite surface proteins Pf12 and Pf41 with Z-scores of 16.9 and 15.4 respectively, followed by gametocyte surface proteins Pfs230 and Pfs48/45 with Z-scores of 15.0 and 12.9, as indicated by a DALI search of the Protein Data Bank (PDB) [51]. All four proteins belong to the 6-cys protein family and share an amino acid sequence identity with Pf12p between 18-27%. The available structures of Pf12 and Pf41 also contain two 6-cys domains whereas structures of Pfs230 and Pfs48/45 feature only a single 6-cys domain (Figure 3B). Structural alignments of Pf12p, Pf12 and Pf41 show that their overall architecture is similar (Figure 3C-E). In all three structures the two 6-cys domains are connected by a short linker and domain-domain contacts are mostly formed between connecting loops of D1 and the five-stranded β-sheet of D2 (Supplementary Figure S3). In the case of Pf12, a surface of 461 Å^2^ is buried between its two domains, 911 Å^2^ for Pf41 and 689 Å^2^ for Pf12p. The two 6-cys domains are tilted against each other in a similar manner, but the relative rotation between D1 and D2 differs in the three structures (Figure 3C).

We have individually aligned the 6-cys domains of Pf12p with the corresponding A- and B-type 6-cys domains of the other family members with known structures (Figure 3D-G). The D1 domain of Pf12p overlays with Pfs230 D1M with a RMSD of 1.8 Å, followed by Pf12 D1 with a RMSD of 2.0 Å and Pf41 D1 with a RMSD of 5.0 Å. The D2 domain of Pf12p overlays with Pfs48/45 6C with a RMSD of 2.2 Å, with Pf12 D2 with a RMSD of 2.4 Å, and with Pf41 D2 with a RMSD of 4.4 Å. The structural alignments show that while the overall domain fold is similar and the spatial position of the cysteine pairs overlay closely, the differences in the length of β-strands, as well as the length and conformation of connecting loops contribute to the relatively high RMSD values.

### Recombinant Pf12p does not interact with Pf41 in solution

Pf12p is a paralog of Pf12 and we wanted to investigate whether the two proteins have redundant roles in the parasite life cycle. Pf41 is a known interaction partner of Pf12 on the merozoite surface and we wanted to determine if Pf12p also had the capability to interact with Pf41. Using BLI we show that Pf41 is able to bind to Pf12 with an equilibrium dissociation constant of K_D_ = 143.7 ± 22.6 nM (Figure 4A). However, for Pf12p, even at the highest concentration of 500 nM we were unable to detect any binding to Pf41 (Figure 4B). Using size exclusion chromatography, we observed complex formation between Pf12 and Pf41 as a higher molecular weight species (Figure 4C). In comparison, there was no indication of a complex forming between Pf12p and Pf41 (Figure 4D). Here, we show that unlike Pf12, Pf12p does not form a complex with Pf41 suggesting that the paralogs may have different functions in the parasite life cycle.

### Pf12p-specific nanobodies bind at the Pf12p D1-D2 domain junction

We used X-ray crystallography to understand how Pf12p-specific nanobodies bind to Pf12p. We purified stable complexes of different nanobodies with either Pf12p D1D2 or Pf12p D2 for crystallization trials. Unfortunately, we obtained no crystallization conditions for Pf12p in complex with B2, C4, or H7, which all bind to the D2 domain of Pf12p and compete with each other. Using Pf12p D1D2, we obtained diffraction quality crystals for Pf12p-B9 and Pf12p-D9 complexes which were determined to 2.0 Å and 3.25 Å, respectively (Figure 5A, B). The Pf12p-B9 structure shows that B9 forms contacts with residues on both Pf12p domains (Figure 5A, C, Table 2). All three CDR loops of B9 are involved in binding Pf12p with an interaction surface of 910 Å^2^ (Figure 5C). The Pf12p-D9 structure also reveals that nanobody D9 forms contacts with residues on both Pf12p domains (Figure 5B, D, Table 2). CDR1 and CDR3 loops of D9 are involved in binding to Pf12p with an interaction surface of 710 Å^2^. The side chains of the CDR2 region of D9 are not well resolved, therefore we are unable to determine the contribution of this CDR loop to binding of Pf12p. While the engagement of the CDR loops with Pf12p differ between B9 and D9, both nanobody epitopes partly overlap, which is consistent with our BLI results showing that B9 and D9 are competing nanobodies.

**Table 1:** Data collection and refinement statistics

|  | Pf12p | Pf12p-B9 | Pf12p-D9 |
| --- | --- | --- | --- |
| **Data collection statistics** |  |  |  |
| Wavelength (Å) | 0.953647 | 0.953725 | 0.953649 |
| Resolution range (Å) | 47.26-2.79<br>(2.96-2.79) | 48.45-2.00<br>(2.12-2.00) | 42.75-3.25<br>(3.44-3.25) |
| Space group | P2<sub>1</sub>2<sub>1</sub>2<sub>1</sub> | P2<sub>1</sub>2<sub>1</sub>2<sub>1</sub> | P2<sub>1</sub>2<sub>1</sub>2<sub>1</sub> |
| Cell axes (Å) (a, b, c) | 52.0, 74.2, 183.9 | 85.0, 107.0, 114.0 | 62.0, 67.1, 111.0 |
| Cell angles (°) ($\alpha$, $\gamma$, $\beta$) | 90, 90, 90 | 90, 90, 90 | 90, 90, 90 |
| Completeness (%) | 99.7 (98.8) | 100.0 (99.9) | 99.8 (99.3) |
| Total no. of reflections | 134842 (20758) | 978968 (159626) | 101019 (15665) |
| Unique reflections | 18315 (2880) | 70873 (11300) | 7710 (1207) |
| Redundancy | 7.3 (7.2) | 13.8 (14.1) | 13.1 (13.0) |
| R<sub>meas</sub> (%) | 19.7 (159.8) | 20.3 (148.7) | 16.0 (210.0) |
| CC<sub>1/2</sub> (%) | 99.5 (71.7) | 99.8 (70.2) | 99.9 (74.1) |
| I/$\sigma$ | 8.44 (1.14) | 10.73 (1.60) | 13.51 (1.44) |
| Wilson B (Å<sup>2</sup>) | 63.12 | 34.60 | 99.23 |
| **Refinement statistics** |  |  |  |
| R<sub>work</sub>/R<sub>free</sub> (%) | 25.0/ 29.5 | 18.0/ 21.7 | 24.5/ 28.6 |
| No. of atoms |  |  |  |
| Protein | 4078 | 6125 | 2756 |
| Water | 10 | 593 | 0 |
| Citrate |  | 72 |  |
| B factors (Å<sup>2</sup>) |  |  |  |
| Chain A | 71.2 | 35.6 | 105.9 |
| Chain B | 67.6 | 35.0 | 112.6 |
| Chain C | 66.2 | 33.4 |  |
| Chain D |  | 37.8 |  |
| Water | 10 | 41.2 | 0 |
| RMDS |  |  |  |
| Bond lengths (Å) | 0.003 | 0.008 | 0.003 |
| Bond angles (°) | 0.625 | 0.932 | 0.578 |
| PDB ID | 7KJ7 | 7KJH | 7KJI |

**Table 2:** Interactions between Pf12p and nanobodies B9 and D9.

Pf12p and nanobody B9 (PDB ID 7KJH)
| Pf12p | Group | Location | Nb B9 | Group | Location | Distance (Å) |
| --- | --- | --- | --- | --- | --- | --- |
| **Hydrogen bonds** |  |  |  |  |  |  |
| Arg 73 | NH2 | D1 | Leu 57 | O | CDR2 | 2.9 |
| Arg 73 | NH1 | D1 | Ala 58 | O | CDR2 | 2.9 |
| Tyr 115 | O | D1 | Phe 103 | N | CDR3 | 2.8 |
| Glu 116 | OE2 | D1 | Gly 102 | N | CDR3 | 2.8 |
| Ile 119 | N | D1 | Tyr 59 | OH | CDR2 | 3.8 |
| Asn 120 | ND2 | D1 | Tyr 59 | OH | CDR2 | 3.6 |
| Asn 269 | O | D2 | Arg 27 | NH1 | CDR1 | 3.2 |
| Asn 269 | OD1 | D2 | Arg 27 | NH1 | CDR1 | 3.0 |
| Lys 273 | NZ | D2 | Gly 26 | O | CDR1 | 3.2 |
| Lys 273 | NZ | D2 | Thr 28 | O | CDR1 | 2.9 |
| Lys 273 | NZ | D2 | Thr 28 | OG1 | CDR1 | 3.5 |
| Tyr 295 | O | D2 | Ser 30 | N | CDR1 | 2.7 |
| Tyr 295 | O | D2 | Ser 30 | OG | CDR1 | 2.8 |
| Gln 297 | O | D2 | Asn 31 | ND2 | CDR1 | 2.9 |
| Gln 297 | OE1 | D2 | Arg 27 | NH2 | CDR1 | 2.9 |
| Gln 297 | OE1 | D2 | Arg 27 | NH1 | CDR1 | 3.0 |
| Gln 297 | N | D2 | Asn 31 | OD1 | CDR1 | 3.3 |
| Lys 299 | NZ | D2 | Gly 104 | O | CDR3 | 3.9 |
| Lys 299 | NZ | D2 | Asn 105 | OD1 | CDR3 | 2.9 |
| **Salt bridges** |  |  |  |  |  |  |
| Glu 116 | OE2 | D1 | Arg 99 | NH1 | CDR3 | 2.9 |
| Glu 116 | OE1 | D1 | Arg 99 | NH2 | CDR3 | 2.9 |
| **Other Pf12p interfacing residues** |  |  |  |  |  |  |
| Val 26 | Phe 69 | Tyr 71 | Leu 72 | Leu 114 | Ser 118 | Asp 121 |
| Asn 122 | Ile 123 | Thr 125 | Asp 127 | Val 128 | Phe 129 | Asp 270 |
| Cys 271 | Phe 272 | Leu 294 | His 296 | Asp 298 | Lys 301 | Thr 304 |

Pf12p and nanobody D9 (PDB ID 7KJI)
| Pf12p | Group | Location | Nb D9 | Group | Location | Distance (Å) |
| --- | --- | --- | --- | --- | --- | --- |
| **Hydrogen bonds** |  |  |  |  |  |  |
| Tyr 71 | OH | D1 | His 32 | NE2 | CDR1 | 3.2 |
| Arg 73 | NH1 | D1 | Lys 75 | O | FR2 | 2.9 |
| Arg 73 | NH1 | D1 | Asn 76 | OD1 | FR2 | 2.8 |
| Glu 116 | OE1 | D1 | Ser 30 | N | CDR1 | 2.8 |
| Glu 116 | OE2 | D1 | Ser 30 | N | CDR1 | 3.0 |
| Asn 120 | ND2 | D1 | Ala 74 | O | FR2 | 2.9 |
| Lys 273 | NZ | D2 | Gly 103 | O | CDR3 | 3.2 |
| Tyr 295 | N | D2 | Thr 104 | OG1 | CDR3 | 3.0 |
| Tyr 295 | O | D2 | Thr 104 | N | CDR3 | 3.8 |
| Tyr 295 | O | D2 | Thr 104 | OG1 | CDR3 | 2.7 |
| Gln 297 | N | D2 | Phe 102 | O | CDR3 | 2.9 |
| **Salt bridges** |  |  |  |  |  |  |
| Lys 273 | NZ | D2 | Asp 100 | OD1 | CDR3 | 3.9 |
| Lys 273 | NZ | D2 | Glu 105 | OE2 | CDR3 | 3.6 |
| **Other Pf12p interfacing residues** |  |  |  |  |  |  |
| Ser 23 | Gly 25 | Val 26 | Asp 28 | Glu 70 | Leu 72 | Ile 75 |
| Leu 114 | Tyr 115 | Ile 123 | Thr 125 | Asp 127 | Val 128 | Phe 129 |
| Ile 293 | Leu 294 | His 296 | Asp 298 | Lys 299 |  |  |

The distance measurements are based on molecules B, D (Pf12p-B9) and molecules A, B (Pf12p-D9).
Interactions and interfacing residues between Pf12p and nanobodies were determined using PISA.

**Fig 5.**
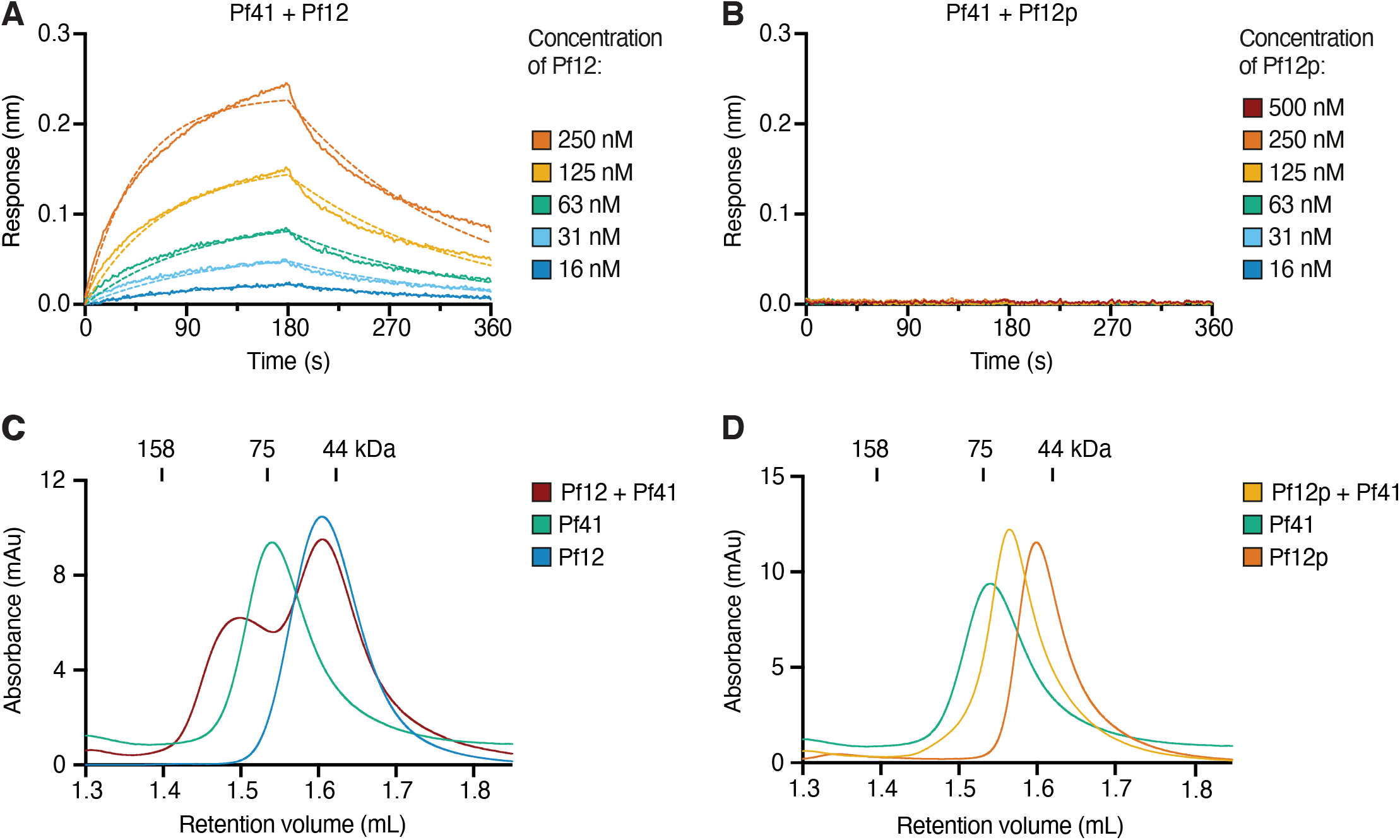
Crystal structures of Pf12p in complex with nanobody B9 and D9, respectively. (**A)** Structure of Pf12p bound to nanobody B9. (**B)** Structure of Pf12p bound to nanobody D9. For panel A and B, the complementary determining regions (CDR) are coloured in light blue (CDR1), blue (CDR2), and dark blue (CDR3). **(C)** Footprint of nanobody B9 on Pf12p D1 and D2 domains. **(D)** Footprint of nanobody D9 on Pf12p D1 and D2 domains. The Pf12p D1 and D2 domains are shown in surface representation in light and dark grey, respectively. The footprint of CDR loops is coloured as described in panel A and B. Coloured Pf12p residues represent those that contact the nanobodies within a distance cutoff of 5 Å. The interaction surface area is indicated. (**E)** Structural alignment of five Pf12p molecules derived from the asymmetric units of the three structures Pf12p (PDB ID 7KJ7), Pf12p bound to nanobody B9 (PDB ID 7KJH) and Pf12p bound to nanobody D9 (PDB ID 7KJI).

To determine if B9 and D9 binding introduces structural changes in Pf12p, we aligned the atomic coordinates of Pf12p obtained from the Pf12p D1D2 structure with that of Pf12p-B9 and Pf12p-D9 (Figure 5E). While only one Pf12p-D9 complex is present in the asymmetric unit of the crystal structure, there are two Pf12p molecules and two Pf12p-B9 complexes in their respective asymmetric units. The five individual chains of Pf12p align with RMSD values of 0.3-0.8 Å. This indicates that there was no major structural change in Pf12p upon binding of the B9 and D9 nanobodies (Figure 5E). Minor positional differences in the β-strand connecting loops between the different Pf12p chains were observed indicating some flexibility in these regions. Our crystal structures of Pf12p-B9 and Pf12p-D9 show that both nanobodies interact with Pf12p at the D1-D2 junction. To our knowledge these are the first available structures of a 6-cys protein containing two 6-cys domains in complex with antibody fragments.

## Discussion

Members of the 6-cys family of proteins are conserved across *Plasmodium* species and play critical roles in parasite invasion, fertilisation, transmission and host immune evasion. However, the precise function of many members remains unknown and structural information is not available for the majority of these surface antigens. In this study, we report the first crystal structure of *P. falciparum* protein Pf12p with its two 6-cys domains. We also characterize a collection of anti-Pf12p nanobodies for their specificity, affinities and their epitope bins. Furthermore, we describe two crystal structures of Pf12p bound to distinct nanobodies, both of which show that nanobodies are able to bind to regions spanning two separate 6-cys domains.

Immunisation of Pf12p in alpacas and subsequent selection of nanobodies using phage display resulted in the identification of ten distinct clonal groups of nanobodies against Pf12p. These ten nanobodies were specific against Pf12p and did not show cross-reactivity towards either Pf12 or Pf41, which are the closest structural homologues to Pf12p. The nanobodies have affinities ranging from ∼3 to 105 nM for binding to Pf12p and their CDR3 regions vary in length between 8 to 21 amino acids. Using BLI, we determined that the antibodies belong to two different epitope bins. One group of five nanobodies bind within domain D2 of Pf12p, but we were unable to obtain crystal structures of this set of nanobodies for detailed epitope determination. In the second group of five nanobodies, binding to Pf12p is partially or completely abrogated in the absence of the D1 domain and we were able to determine the structure of two Pf12p-nanobody complexes of this group of antibodies.

We observed that the two Pf12p-specific nanobodies interact with the interdomain region of Pf12p and simultaneously engage both domains of this protein. This mode of binding is novel for antibodies that recognize 6-cys protein family members. Published crystal structures of inhibitory antibodies are all located within single 6-cys domains of Pfs230 and Pfs48/45. Both anti-Pfs230 antibody 4F12 and anti-Pfs48/45 antibody 85RF45.1 bind their respective antigen at an edge of the β-sandwich of domain D1 and D3, respectively, by engaging many residues of the β-strand connecting loops. By comparison, our nanobodies B9 and D9 form most interactions with residues located on β-sheet A of each domain. We propose that nanobodies are able to bind to the interdomain regions and provide a tool for identifying novel inhibitory epitopes of the 6-cys protein family.

In the three available structures of containing two 6-cys domains in Pf12, Pf41 and Pf12p, D1 and D2 are rotated against each other in a similar manner and interdomain interactions bury extended surface areas from solvent exposure (461-911 Å^2^). The smallest D1-D2 interface is present in the structure of Pf12, but here 33 residues are missing in D1 which could (partly) contribute to further interdomain interactions. While the described D1-D2 domain interactions of Pf12 are of hydrophobic nature and predicted to allow mobility between D1 and D2 [26], interdomain contacts of Pf41 and Pf12p involve hydrogen bonds as well as hydrophobic and aromatic interactions suggesting that the domain arrangement of 6-cys protein tandems of A-type and B-type are rigid rather than flexible. In comparison, the tandem domains of the related SRS protein family in *T. gondii*, are organized in a linear head-to tail arrangement forming no or limited interdomain interactions [31-33]. The two domains of this protein family are therefore flexible towards each other and the domain-connecting linker is believed to facilitate structural adaptations during ligand binding [31, 32, 52]. The predicted ligand binding site of parallel orientated SRS-homodimers are located at the dimer interface of the D1 domains [31, 33].

Length and size of β-strand connecting loops vary between 6-cys protein family members. Our crystal structures show that Pf12p has a 40 residue-long loop in the D2 domain, which is disordered and asparagine-rich. The extreme AT-rich genome of *P. falciparum* features an abundance of trinucleotide repeats coding for asparagine that causes a wealth of low-complexity, asparagine-rich regions in *P. falciparum* proteins, such as Pf12p. About 30% of the *P. falciparum* proteome contains such low-complexity amino acid repeats with stretches of 37 residues on average [53]. The repeats are found in all protein families and in every stage of the life cycle. For the 6-cys family, asparagine-rich regions with six or more asparagine residues in a row, are present in Pf12p, Pf52, PfLISP2 and PfB9. In other *Plasmodium* species asparagine-rich regions are rare, except in *P. reichenowi* and this repeat region is not conserved in P12p orthologs across *Plasmodium* species. The functional role of these asparagine-rich regions will need to be determined.

Gene duplication played a role in the expansion of the 6-cys protein family resulting in four paralogous pairs of genes [30, 36, 54, 55]. The family members Pfs230 and Pfs230p are paralogs, as are Pfs48/45 and Pf47, Pf36 and Pf52, and Pf12 and Pf12p. The amino acid sequence identity is low among the pairs of paralogs. Pf12p is reported to be transcribed in blood stages of infection and present in sporozoites but the protein is not associated with any function to date [27, 41, 42]. Pf12 is present on the surface of schizonts and merozoites [25, 26]. The biological role of Pf12 is unknown, but its ability to interact with Pf41 is well characterized [24-26]. In comparison to Pf12, the Pf12p protein shows no evidence of interacting with Pf41 suggesting that the roles of Pf12 and Pf12p are probably not interchangeable.

The 6-cys family of proteins are an important class of surface-proteins involved in different functions of the *Plasmodium* parasite life cycle. In conclusion, we have generated and characterized ten different nanobodies that bind Pf12p with high affinity and specificity. Crystal structures of two nanobody-Pf12p complexes reveal a novel binding mode of antibodies that recognize 6-cys proteins by engaging the interdomain region of the protein. We propose that nanobodies targeting 6-cys proteins are a useful tool to identify new inhibitory epitopes for 6-cys proteins and will contribute to unravelling the diverse functions of this protein family.

## Data availability

Coordinates and structure factors have been deposited in the Protein Data Bank (PDB) under PDB ID 7KJ7 for Pf12p, 7KJH for Pf12p-B9 complex and 7KJI for Pf12p-D9 complex.

## Acknowledgement

We thank Janet Newman and Bevan Marshall from the CSIRO Collaborative Crystallization Centre (CSIRO; Parkville, Australia) for assistance with setting up the crystallization screens. This research was undertaken using the MX2 beamline at the Australian Synchrotron and we thank the MX2 beamline staff at the Australian Synchrotron for their assistance during data collection. W.-H.T. is a Howard Hughes Medical Institute-Wellcome Trust International Research Scholar (208693/Z/17/Z) and supported by National Health and Medical Research Council of Australia (GNT1143187, GNT1160042, GNT1160042, GNT1154937).

## Author contribution statement

M.H.D. expressed and purified all recombinant proteins and prepared samples for protein crystallography, crystallized Pf12p and Pf12p-Nb complexes, collected diffraction data and determined the crystal structures. L.-J.C. performed bio-layer interferometry measurements for affinities and epitope competition. A.A. generated the nanobody phage library, performed the bio-panning, nanobody sequencing and nanobody specificity ELISAs. P.P. performed complex formation studies using size exclusion chromatography. Recombinant nanobodies were purified with assistance from A.A, S.K., AWL, and Y-C.C. W.-H.T. and M.H.D. conceived the project, designed the experiments and analysed the data. All authors assisted in manuscript preparation.

## Figure legends

**Supplementary Fig S1. BLI-affinity measurements with immobilized nanobodies and Pf12p D1D2 in solution. (A)** Representative binding curves of six different Pf12p D1D2 concentrations to immobilized nanobodies are shown and were fitted to a 1:1 binding model. Corresponding K_D_ values are indicated. **(B)** Table containing determined kinetic and affinity data from three independent experiments showing the mean and standard error of the mean (SEM).

**Supplementary Fig S2. Amino acid sequence alignment of Pf12p with orthologs of different *Plasmodium species***. The program ClustalO was used for the alignment [56]. The *Plasmodium* species with uniprot ID of the corresponding P12p protein is indicated on the left hand side of the alignment. Conserved cysteine residues are highlighted in yellow and the asparagine-rich region of Pf12p is highlighted in purple.

**Supplementary Fig S3. Interdomain interactions of Pf12 D1-D2, Pf41 D1-D2 and Pf12p D1-D2**. Interdomain linker regions are highlighted in green and sidechain residues at the interface of the two domains, D1 and D2, are shown in ball and stick representation. In all three structures the domain-domain contacts are mostly formed between connecting loops of D1 and the five-stranded β-sheet of D2. **(A)** Pf12, **(B)** Pf41, **(C)** Pf12p.

## References

1 WHO. (2019) World Malaria Report: 2019. ed.)^eds.)

2 Aurrecoechea, C., Brestelli, J., Brunk, B. P., Dommer, J., Fischer, S., Gajria, B., Gao, X., Gingle, A., Grant, G., Harb, O. S., Heiges, M., Innamorato, F., Iodice, J., Kissinger, J. C., Kraemer, E., Li, W., Miller, J. A., Nayak, V., Pennington, C., Pinney, D. F., Roos, D. S., Ross, C., Stoeckert, C. J., Jr., Treatman, C. and Wang, H. (2009) PlasmoDB: a functional genomic database for malaria parasites. Nucleic Acids Res. 37, D539–543

3 Arredondo, S. A., Swearingen, K. E., Martinson, T., Steel, R., Dankwa, D. A., Harupa, A., Camargo, N., Betz, W., Vigdorovich, V., Oliver, B. G., Kangwanrangsan, N., Ishino, T., Sather, N., Mikolajczak, S., Vaughan, A. M., Torii, M., Moritz, R. L. and Kappe, S. H. I. (2018) The Micronemal Plasmodium Proteins P36 and P52 Act in Concert to Establish the Replication-Permissive Compartment Within Infected Hepatocytes. Front Cell Infect Microbiol. 8, 413

4 Garcia, J., Curtidor, H., Pinzon, C. G., Vanegas, M., Moreno, A. and Patarroyo, M. E. (2009) Identification of conserved erythrocyte binding regions in members of the Plasmodium falciparum Cys6 lipid raft-associated protein family. Vaccine. 27, 3953–3962

5 Kennedy, A. T., Schmidt, C. Q., Thompson, J. K., Weiss, G. E., Taechalertpaisarn, T., Gilson, P. R., Barlow, P. N., Crabb, B. S., Cowman, A. F. and Tham, W. H. (2016) Recruitment of Factor H as a Novel Complement Evasion Strategy for Blood-Stage Plasmodium falciparum Infection. J Immunol. 196, 1239–1248

6 Molina-Cruz, A., Canepa, G. E., Alves, E. S. T. L., Williams, A. E., Nagyal, S., Yenkoidiok-Douti, L., Nagata, B. M., Calvo, E., Andersen, J., Boulanger, M. J. and Barillas-Mury, C. (2020) Plasmodium falciparum evades immunity of anopheline mosquitoes by interacting with a Pfs47 midgut receptor. Proc Natl Acad Sci U S A. 117, 2597–2605

7 Ramiro, R. S., Khan, S. M., Franke-Fayard, B., Janse, C. J., Obbard, D. J. and Reece, S. E. (2015) Hybridization and pre-zygotic reproductive barriers in Plasmodium. Proc Biol Sci. 282, 20143027

8 van Dijk, M. R., Janse, C. J., Thompson, J., Waters, A. P., Braks, J. A., Dodemont, H. J., Stunnenberg, H. G., van Gemert, G. J., Sauerwein, R. W. and Eling, W. (2001) A central role for P48/45 in malaria parasite male gamete fertility. Cell. 104, 153–164

9 Theisen, M., Jore, M. M. and Sauerwein, R. (2017) Towards clinical development of a Pfs48/45-based transmission blocking malaria vaccine. Expert Rev Vaccines. 16, 329–336

10 Williamson, K. C. (2003) Pfs230: from malaria transmission-blocking vaccine candidate toward function. Parasite Immunol. 25, 351–359

11 Eksi, S., Czesny, B., van Gemert, G. J., Sauerwein, R. W., Eling, W. and Williamson, K. C. (2006) Malaria transmission-blocking antigen, Pfs230, mediates human red blood cell binding to exflagellating male parasites and oocyst production. Mol Microbiol. 61, 991–998

12 Kumar, N. (1987) Target antigens of malaria transmission blocking immunity exist as a stable membrane bound complex. Parasite Immunol. 9, 321–335

13 Kumar, N. and Wizel, B. (1992) Further characterization of interactions between gamete surface antigens of Plasmodium falciparum. Mol Biochem Parasitol. 53, 113–120

14 Carter, R., Graves, P. M., Keister, D. B. and Quakyi, I. A. (1990) Properties of epitopes of Pfs 48/45, a target of transmission blocking monoclonal antibodies, on gametes of different isolates of Plasmodium falciparum. Parasite Immunol. 12, 587–603

15 Foo, A., Carter, R., Lambros, C., Graves, P., Quakyi, I., Targett, G. A., Ponnudurai, T. and Lewis, G. E., Jr. (1991) Conserved and variant epitopes of target antigens of transmission-blocking antibodies among isolates of Plasmodium falciparum from Malaysia. Am J Trop Med Hyg. 44, 623–631

16 Molina-Cruz, A., Garver, L. S., Alabaster, A., Bangiolo, L., Haile, A., Winikor, J., Ortega, C., van Schaijk, B. C., Sauerwein, R. W., Taylor-Salmon, E. and Barillas-Mury, C. (2013) The human malaria parasite Pfs47 gene mediates evasion of the mosquito immune system. Science. 340, 984–987

17 van Dijk, M. R., van Schaijk, B. C., Khan, S. M., van Dooren, M. W., Ramesar, J., Kaczanowski, S., van Gemert, G. J., Kroeze, H., Stunnenberg, H. G., Eling, W. M., Sauerwein, R. W., Waters, A. P. and Janse, C. J. (2010) Three members of the 6-cys protein family of Plasmodium play a role in gamete fertility. PLoS Pathog. 6, e1000853

18 van Schaijk, B. C., van Dijk, M. R., van de Vegte-Bolmer, M., van Gemert, G. J., van Dooren, M. W., Eksi, S., Roeffen, W. F., Janse, C. J., Waters, A. P. and Sauerwein, R. W. (2006) Pfs47, paralog of the male fertility factor Pfs48/45, is a female specific surface protein in Plasmodium falciparum. Mol Biochem Parasitol. 149, 216–222

19 Kaushansky, A., Douglass, A. N., Arang, N., Vigdorovich, V., Dambrauskas, N., Kain, H. S., Austin, L. S., Sather, D. N. and Kappe, S. H. (2015) Malaria parasites target the hepatocyte receptor EphA2 for successful host infection. Science. 350, 1089–1092

20 Manzoni, G., Marinach, C., Topcu, S., Briquet, S., Grand, M., Tolle, M., Gransagne, M., Lescar, J., Andolina, C., Franetich, J. F., Zeisel, M. B., Huby, T., Rubinstein, E., Snounou, G., Mazier, D., Nosten, F., Baumert, T. F. and Silvie, O. (2017) Plasmodium P36 determines host cell receptor usage during sporozoite invasion. Elife. 6

21 Osier, F. H., Mackinnon, M. J., Crosnier, C., Fegan, G., Kamuyu, G., Wanaguru, M., Ogada, E., McDade, B., Rayner, J. C., Wright, G. J. and Marsh, K. (2014) New antigens for a multicomponent blood-stage malaria vaccine. Sci Transl Med. 6, 247ra102

22 Richards, J. S., Arumugam, T. U., Reiling, L., Healer, J., Hodder, A. N., Fowkes, F. J., Cross, N., Langer, C., Takeo, S., Uboldi, A. D., Thompson, J. K., Gilson, P. R., Coppel, R. L., Siba, P. M., King, C. L., Torii, M., Chitnis, C. E., Narum, D. L., Mueller, I., Crabb, B. S., Cowman, A. F., Tsuboi, T. and Beeson, J. G. (2013) Identification and prioritization of merozoite antigens as targets of protective human immunity to Plasmodium falciparum malaria for vaccine and biomarker development. J Immunol. 191, 795–809

23 Sanders, P. R., Gilson, P. R., Cantin, G. T., Greenbaum, D. C., Nebl, T., Carucci, D. J., McConville, M. J., Schofield, L., Hodder, A. N., Yates, J. R., 3rd and Crabb, B. S. (2005) Distinct protein classes including novel merozoite surface antigens in Raft-like membranes of Plasmodium falciparum. J Biol Chem. 280, 40169–40176

24 Parker, M. L., Peng, F. and Boulanger, M. J. (2015) The Structure of Plasmodium falciparum Blood-Stage 6-Cys Protein Pf41 Reveals an Unexpected Intra-Domain Insertion Required for Pf12 Coordination. PLoS One. 10, e0139407

25 Taechalertpaisarn, T., Crosnier, C., Bartholdson, S. J., Hodder, A. N., Thompson, J., Bustamante, L. Y., Wilson, D. W., Sanders, P. R., Wright, G. J., Rayner, J. C., Cowman, A. F., Gilson, P. R. and Crabb, B. S. (2012) Biochemical and functional analysis of two Plasmodium falciparum blood-stage 6-cys proteins: P12 and P41. PLoS One. 7, e41937

26 Tonkin, M. L., Arredondo, S. A., Loveless, B. C., Serpa, J. J., Makepeace, K. A., Sundar, N., Petrotchenko, E. V., Miller, L. H., Grigg, M. E. and Boulanger, M. J. (2013) Structural and biochemical characterization of Plasmodium falciparum 12 (Pf12) reveals a unique interdomain organization and the potential for an antiparallel arrangement with Pf41. J Biol Chem. 288, 12805–12817

27 Gilson, P. R., Nebl, T., Vukcevic, D., Moritz, R. L., Sargeant, T., Speed, T. P., Schofield, L. and Crabb, B. S. (2006) Identification and stoichiometry of glycosylphosphatidylinositol-anchored membrane proteins of the human malaria parasite Plasmodium falciparum. Mol Cell Proteomics. 5, 1286–1299

28 Arredondo, S. A., Cai, M., Takayama, Y., MacDonald, N. J., Anderson, D. E., Aravind, L., Clore, G. M. and Miller, L. H. (2012) Structure of the Plasmodium 6-cysteine s48/45 domain. Proc Natl Acad Sci U S A. 109, 6692–6697

29 Carter, R., Coulson, A., Bhatti, S., Taylor, B. J. and Elliott, J. F. (1995) Predicted disulfide-bonded structures for three uniquely related proteins of Plasmodium falciparum, Pfs230, Pfs48/45 and Pf12. Mol Biochem Parasitol. 71, 203–210

30 Gerloff, D. L., Creasey, A., Maslau, S. and Carter, R. (2005) Structural models for the protein family characterized by gamete surface protein Pfs230 of Plasmodium falciparum. Proc Natl Acad Sci U S A. 102, 13598–13603

31 Crawford, J., Grujic, O., Bruic, E., Czjzek, M., Grigg, M. E. and Boulanger, M. J. (2009) Structural characterization of the bradyzoite surface antigen (BSR4) from Toxoplasma gondii, a unique addition to the surface antigen glycoprotein 1-related superfamily. J Biol Chem. 284, 9192–9198

32 Crawford, J., Lamb, E., Wasmuth, J., Grujic, O., Grigg, M. E. and Boulanger, M. J. (2010) Structural and functional characterization of SporoSAG: a SAG2-related surface antigen from Toxoplasma gondii. J Biol Chem. 285, 12063–12070

33 He, X. L., Grigg, M. E., Boothroyd, J. C. and Garcia, K. C. (2002) Structure of the immunodominant surface antigen from the Toxoplasma gondii SRS superfamily. Nat Struct Biol. 9, 606–611

34 Annoura, T., van Schaijk, B. C., Ploemen, I. H., Sajid, M., Lin, J. W., Vos, M. W., Dinmohamed, A. G., Inaoka, D. K., Rijpma, S. R., van Gemert, G. J., Chevalley-Maurel, S., Kielbasa, S. M., Scheltinga, F., Franke-Fayard, B., Klop, O., Hermsen, C. C., Kita, K., Gego, A., Franetich, J. F., Mazier, D., Hoffman, S. L., Janse, C. J., Sauerwein, R. W. and Khan, S. M. (2014) Two Plasmodium 6-Cys family-related proteins have distinct and critical roles in liver-stage development. FASEB J. 28, 2158–2170

35 Manger, I. D., Hehl, A. B. and Boothroyd, J. C. (1998) The surface of Toxoplasma tachyzoites is dominated by a family of glycosylphosphatidylinositol-anchored antigens related to SAG1. Infect Immun. 66, 2237–2244

36 Templeton, T. J. and Kaslow, D. C. (1999) Identification of additional members define a Plasmodium falciparum gene superfamily which includes Pfs48/45 and Pfs230. Mol Biochem Parasitol. 101, 223–227

37 Wasmuth, J. D., Pszenny, V., Haile, S., Jansen, E. M., Gast, A. T., Sher, A., Boyle, J. P., Boulanger, M. J., Parkinson, J. and Grigg, M. E. (2012) Integrated bioinformatic and targeted deletion analyses of the SRS gene superfamily identify SRS29C as a negative regulator of Toxoplasma virulence. mBio. 3

38 Kundu, P., Semesi, A., Jore, M. M., Morin, M. J., Price, V. L., Liang, A., Li, J., Miura, K., Sauerwein, R. W., King, C. R. and Julien, J. P. (2018) Structural delineation of potent transmission-blocking epitope I on malaria antigen Pfs48/45. Nat Commun. 9, 4458

39 Lennartz, F., Brod, F., Dabbs, R., Miura, K., Mekhaiel, D., Marini, A., Jore, M. M., Sogaard, M. M., Jorgensen, T., de Jongh, W. A., Sauerwein, R. W., Long, C. A., Biswas, S. and Higgins, M. K. (2018) Structural basis for recognition of the malaria vaccine candidate Pfs48/45 by a transmission blocking antibody. Nat Commun. 9, 3822

40 Singh, K., Burkhardt, M., Nakuchima, S., Herrera, R., Muratova, O., Gittis, A. G., Kelnhofer, E., Reiter, K., Smelkinson, M., Veltri, D., Swihart, B. J., Shimp, R., Jr., Nguyen, V., Zhang, B., MacDonald, N. J., Duffy, P. E., Garboczi, D. N. and Narum, D. L. (2020) Structure and function of a malaria transmission blocking vaccine targeting Pfs230 and Pfs230-Pfs48/45 proteins. Commun Biol. 3, 395

41 Lasonder, E., Janse, C. J., van Gemert, G. J., Mair, G. R., Vermunt, A. M., Douradinha, B. G., van Noort, V., Huynen, M. A., Luty, A. J., Kroeze, H., Khan, S. M., Sauerwein, R. W., Waters, A. P., Mann, M. and Stunnenberg, H. G. (2008) Proteomic profiling of Plasmodium sporozoite maturation identifies new proteins essential for parasite development and infectivity. PLoS Pathog. 4, e1000195

42 Lindner, S. E., Swearingen, K. E., Harupa, A., Vaughan, A. M., Sinnis, P., Moritz, R. L. and Kappe, S. H. (2013) Total and putative surface proteomics of malaria parasite salivary gland sporozoites. Mol Cell Proteomics. 12, 1127–1143

43 Pardon, E., Laeremans, T., Triest, S., Rasmussen, S. G., Wohlkonig, A., Ruf, A., Muyldermans, S., Hol, W. G., Kobilka, B. K. and Steyaert, J. (2014) A general protocol for the generation of Nanobodies for structural biology. Nat Protoc. 9, 674–693

44 Kabsch, W. (2010) Xds. Acta Crystallogr D Biol Crystallogr. 66, 125–132

45 McCoy, A. J., Grosse-Kunstleve, R. W., Adams, P. D., Winn, M. D., Storoni, L. C. and Read, R. J. (2007) Phaser crystallographic software. J Appl Crystallogr. 40, 658–674

46 Emsley, P., Lohkamp, B., Scott, W. G. and Cowtan, K. (2010) Features and development of Coot. Acta Crystallogr D Biol Crystallogr. 66, 486–501

47 Adams, P. D., Grosse-Kunstleve, R. W., Hung, L. W., Ioerger, T. R., McCoy, A. J., Moriarty, N. W., Read, R. J., Sacchettini, J. C., Sauter, N. K. and Terwilliger, T. C. (2002) PHENIX: building new software for automated crystallographic structure determination. Acta Crystallogr D Biol Crystallogr. 58, 1948–1954

48 Afonine, P. V., Grosse-Kunstleve, R. W., Echols, N., Headd, J. J., Moriarty, N. W., Mustyakimov, M., Terwilliger, T. C., Urzhumtsev, A., Zwart, P. H. and Adams, P. D. (2012) Towards automated crystallographic structure refinement with phenix.refine. Acta Crystallogr D Biol Crystallogr. 68, 352–367

49 The PyMOL Molecular Graphics System, Version 2.3.0, Schrödinger, LLC. ed.)^eds.)

50 Krissinel, E. (2015) Stock-based detection of protein oligomeric states in jsPISA. Nucleic Acids Res. 43, W314–319

51 Holm, L. (2020) DALI and the persistence of protein shape. Protein Sci. 29, 128–140

52 Graille, M., Stura, E. A., Bossus, M., Muller, B. H., Letourneur, O., Battail-Poirot, N., Sibai, G., Gauthier, M., Rolland, D., Le Du, M. H. and Ducancel, F. (2005) Crystal structure of the complex between the monomeric form of Toxoplasma gondii surface antigen 1 (SAG1) and a monoclonal antibody that mimics the human immune response. J Mol Biol. 354, 447–458

53 Muralidharan, V. and Goldberg, D. E. (2013) Asparagine repeats in Plasmodium falciparum proteins: good for nothing? PLoS Pathog. 9, e1003488

54 Gardner, M. J., Tettelin, H., Carucci, D. J., Cummings, L. M., Aravind, L., Koonin, E. V., Shallom, S., Mason, T., Yu, K., Fujii, C., Pederson, J., Shen, K., Jing, J., Aston, C., Lai, Z., Schwartz, D. C., Pertea, M., Salzberg, S., Zhou, L., Sutton, G. G., Clayton, R., White, O., Smith, H. O., Fraser, C. M., Adams, M. D., Venter, J. C. and Hoffman, S. L. (1998) Chromosome 2 sequence of the human malaria parasite Plasmodium falciparum. Science. 282, 1126–1132

55 Thompson, J., Janse, C. J. and Waters, A. P. (2001) Comparative genomics in Plasmodium: a tool for the identification of genes and functional analysis. Mol Biochem Parasitol. 118, 147–154

56 Madeira, F., Park, Y. M., Lee, J., Buso, N., Gur, T., Madhusoodanan, N., Basutkar, P., Tivey, A. R. N., Potter, S. C., Finn, R. D. and Lopez, R. (2019) The EMBL-EBI search and sequence analysis tools APIs in 2019. Nucleic Acids Res. 47, W636–W641

