## Supplemental Figures for "Nanobody generation and structural characterization of *Plasmodium falciparum* 6-cysteine protein Pf12p"

### Figure S1

**A**

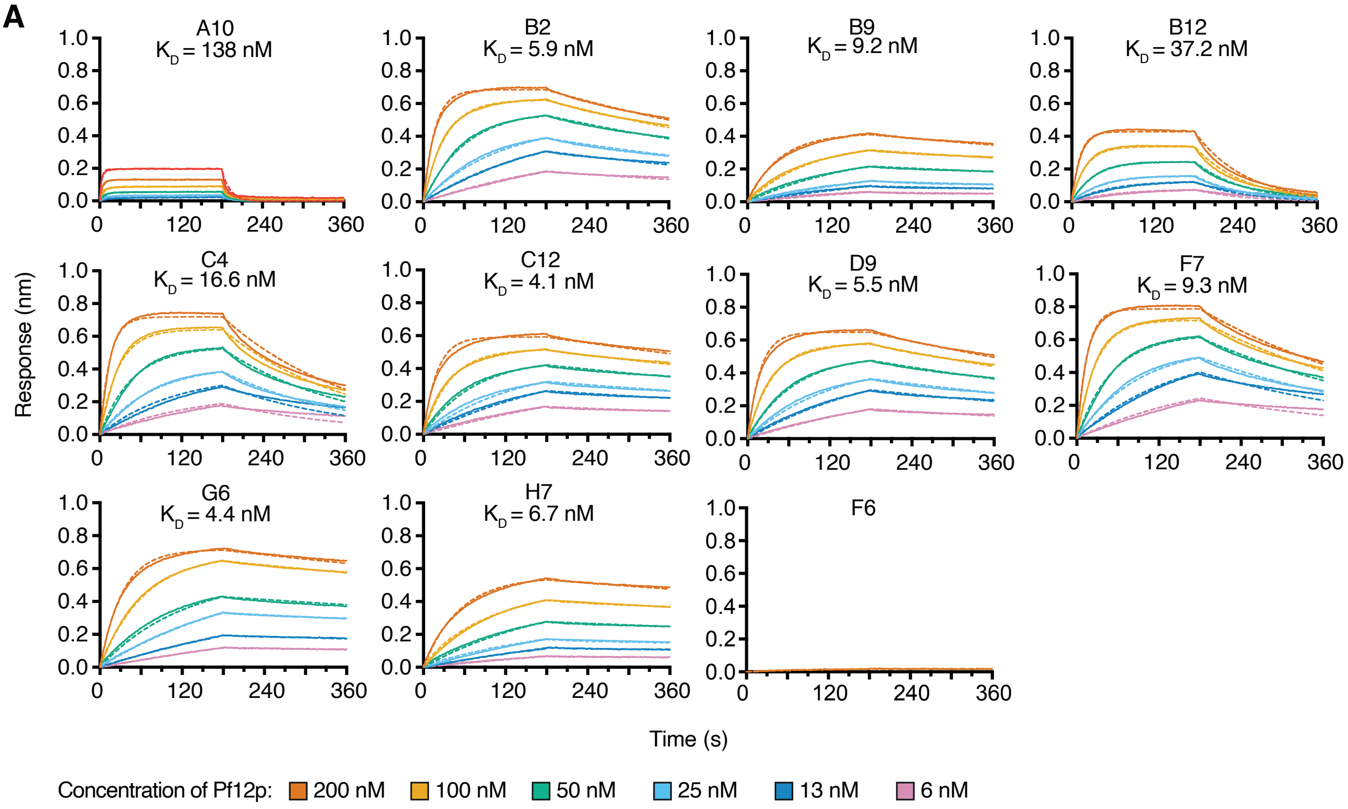


**B**

| mAb | $K_D$ (nM) | $k_a$ (x $10^5$ $M^{-1}s^{-1}$) | $k_d$ (x $10^{-3}$ $s^{-1}$) |
|---|---|---|---|
| A10 | 105.10 (± 24.89) | 24.89 (± 7.46) | 7.46 (± 0.43) |
| B2 | 6.76 (± 0.75) | 0.75 (± 3.00) | 3.00 (± 0.57) |
| B9 | 7.48 (± 0.88) | 0.88 (± 1.21) | 1.21 (± 0.27) |
| B12 | 25.27 (± 5.67) | 5.67 (± 5.38) | 5.38 (± 0.66) |
| C4 | 17.80 (± 2.73) | 2.73 (± 4.62) | 4.62 (± 1.73) |
| C12 | 3.96 (± 0.77) | 0.77 (± 2.68) | 2.68 (± 0.37) |
| D9 | 5.19 (± 0.98) | 0.98 (± 2.96) | 2.96 (± 0.44) |
| F7 | 10.63 (± 1.33) | 1.33 (± 3.81) | 3.81 (± 0.88) |
| G6 | 3.22 (± 0.81) | 0.81 (± 2.23) | 2.23 (± 0.67) |
| H7 | 6.71 (± 0.50) | 0.50 (± 1.12) | 1.12 (± 0.23) |

### Figure S2

```
P.falciparum_C6KSX1      -MHIVSFII-----------FFFA-----------------------LFFPISICYKIN 24
P.berghei_Q4Z5T0         -----------------------------------------------------MIQNI 5
P.knowlesi_A0A1Y3DHK6    -MRVRHFSFLRLFLLLSLLVYHLPVQKQRHRSIPSWKYPDE-GDDHPPQNAFSHMANQVK 58
P.reichenowi_A0A151LP59  -MHIVSFLI-----------FFFA-----------------------LFFPISICYKIN 24
P.malariae_A0A1D3SN11    -MQLVIFAI---------------VQ----------------------FLYFFSFAYKTN 22
P.ovale_A0A1A8VY82       -MPVVKSEG---------------TQ-----------------------ESSRERVQKSK 21
P.chabaudi_A0A4V0K2Q1    MMKTYFWLA---------------VH-----------------------FFSSFWMIQNI 22
P.yoelii_A0A078K5Q3      MMRIYFWLA---------------MH-----------------------IFSSFWMIQNI 22
P.vivax_A0A1G4H080       -MRVGHFSFVRLLLLQSLLVQRLPVQKQQQSSIPARKHPEEEGGGPSPQNALAQVANQMK 59
                                                                             :

P.falciparum_C6KSX1      GVCDFSSEGLSLLPEEKLD---------FSVSRNVDKLSDENNVRHCVHFSKGFEYLRFI 75
P.berghei_Q4Z5T0         EICDFSRGSLDVALMNNKILIDN--N-------LKEENYNDNNIKHCVIFTKGLEIFTFI 56
P.knowlesi_A0A1Y3DHK6    GICDFSRGPLNVSTTENEIVPLLVQAELHAGSAPLSDAHTDEPVQRCVQFTKGMEVLTFV 118
P.reichenowi_A0A151LP59  GVCDFSSEGLSLLPEEKLD---------FSASRNVDKLSDENNVRHCVHFSKGFEYLRFI 75
P.malariae_A0A1D3SN11    DDCDFTREPLNVAWNNRNNALAPVDM-------EDEPYDNDNNIKYCVKFTKGFEILTFI 75
P.ovale_A0A1A8VY82       GECDFTRGSLDVSRSGNNKVAL---F-------EREEEAYDSNVKHCVKFTKGLEIMTFI 71
P.chabaudi_A0A4V0K2Q1    EICDFSKESLDVALTKDKSVIGN--S-------SNEENHSDNNIKHCVKFTKGFEIFTFI 73
P.yoelii_A0A078K5Q3      EICDFSRDSLDVTLMNNKIVIDN--N-------LKEENYNDNNIKHCVKFTKGLEIFTFI 73
P.vivax_A0A1G4H080       GTCDFSRAPLNVSCSENEIVALPGGEGAM-VSGTTGSTANDERARHCVQFTKGFDVLTFV 118
                           ***:   *.:                                :.  : ** *:**:: : *:

P.falciparum_C6KSX1      CPMRK--DNYEGIEIRPVECFEYIHI-EGREHKLSEILKGSLYEKSINDNIMTRDVFIPP 132
P.berghei_Q4Z5T0         CPKGNNNDNYKGVEIRPEQCFEKVRI-NGKEENLKDILKGVVIEKKETDTEIIRKALIPP 115
P.knowlesi_A0A1Y3DHK6    CPKRNT-EDYIGVEIRPMECFEKVRMHNGNKKKLNNVLKGVQLENIDTDSLSIRKVFIPP 177
P.reichenowi_A0A151LP59  CPMRK--DNYEGIEIRPVECFEFVRI-EGREQKLGEILKGSLYEKSINDNIMTRDVFIPP 132
P.malariae_A0A1D3SN11    CPKKG--INYEGIEMRPKECFEKIRI-NGRDENFTEILKGSIFESSETDSIIIRRVFIPP 132
P.ovale_A0A1A8VY82       CPKRS--IHYEAIEIRPTDCFVKVRV-NGIEERLNDFMKGVIFESRENDLSNIRKVFIPP 128
P.chabaudi_A0A4V0K2Q1    CPKGNNNDNYNGIEIRPVQCFEKVRI-NGKEENLKDVLKGVITENKETDTSIIRKAFIPP 132
P.yoelii_A0A078K5Q3      CPKGNNNDNYKGIEIRPEQCFEKVRI-NGKEENLKDILKGVIIEKKETDTEIIRKAFIPP 132
P.vivax_A0A1G4H080       CPKRSS-EDYSGVEIRPMSCFETVRRTDGTNQQLSEVLKGVQLENRDTDLLSIRRVFIPP 177
                         **       .* .:*:** .**  ::  :*  ...: :.:**   *.   .*    *  .:***

P.falciparum_C6KSX1      TIYEDMFFECTCDNSLTFKNNMIGIRGIMKIHLKKNILYGCDFDHDEKL----------- 181
P.berghei_Q4Z5T0         TIYQDMSFECSCDNSLTIKDNYIGARGIMKVHLKKNIIFGCDFNYDSNE----------- 164
P.knowlesi_A0A1Y3DHK6    TIYRNIIFECTCDNSLSFWNNKMGTRGIMRVHLRKNIVFGCDFDHRGGRENILEVEGELP 237
P.reichenowi_A0A151LP59  TIYEDMFFECTCDNSLTFKNNMIGIKGIMKIHLKKNILYGCDFDHDEKL----------- 181
P.malariae_A0A1D3SN11    TIYADMVIECTCDNSLTFKENFIGARGIMRVHLKRNKIFGCDFDSNIDGDDS-------- 184
P.ovale_A0A1A8VY82       TIYEDIVFECTCDNSLTFGDNQIGTRGIMRVHLKKNLVFGCDFDYDVMN----------- 177
P.chabaudi_A0A4V0K2Q1    TIYNDMSFECSCDNSLTIKDNTIGARGIMRVHLKKNKIFGCDFNYDASD----------- 181
P.yoelii_A0A078K5Q3      TIYQDMSFECSCDNSLTIKDNYIGARGIMKVHLKKNIIFGCDFNYDTNE----------- 181
P.vivax_A0A1G4H080       TIYQNFIFECSCDNSLTFWKNKMGARGIMRVHLRRNILFGCDFDHTGGVEYAGGLGGELP 237
                         *** :: :**:*****:: .* :* :***::**::* ::****:

P.falciparum_C6KSX1      -------------------------------------MKNKTAFTNFYDKQKILPLIGNN 204
P.berghei_Q4Z5T0         ----------------------------------PKLSNGKSAFAQFYDKQV-------- 182
P.knowlesi_A0A1Y3DHK6    AVDEAN----------RNTTGDWAFWRNAGPSAEELAERNKTAFSQFYTSEEV------- 280
P.reichenowi_A0A151LP59  -------------------------------------MKNKTAFTNFYDKQKILSLIDNN 204
P.malariae_A0A1D3SN11    -----TYG----RSDGGTRN----KCEGRLDNNCEESKSGRSAFVKYYDKSK-------- 223
P.ovale_A0A1A8VY82       -------------------------------------SRKRSAFVSFYEKNE-------- 192
P.chabaudi_A0A4V0K2Q1    ----------------------------------TKFSNGKSAFTNFYDNQA-------- 199
P.yoelii_A0A078K5Q3      ----------------------------------PKHSNGKSAFARFYDKKII------- 200
P.vivax_A0A1G4H080       MADEASVGSAEDRSAGGGSADDWAFWRNAGPSAEELAEKNKTAFTNFYPPGEV------- 290
                                                                 ::**   :*

P.falciparum_C6KSX1      NNDDDNNDDDNNNDNNNNDNNNNNNNNNNNNNNNNNNNITCNVTIKKSQVYLGIICPDGY 264
P.berghei_Q4Z5T0         -------------------------------V-DSNKNIICNTQVNNKEVYLGLVCPEGY 210
P.knowlesi_A0A1Y3DHK6    -------------------------------NDAKDKGIICNVKITKREVYLGLVCPSGY 309
P.reichenowi_A0A151LP59  NNNNNDDDDD--------------------NNNNNNNNITCNVTIKQSQVYLGIICPDGY 244
P.malariae_A0A1D3SN11    -------------------------------I-RSNESITCNVTINKKEVYLGLICPEGY 251
P.ovale_A0A1A8VY82       -------------------------------I-KPGEEIVCNVKITKKEVYLGLICPEGY 220
P.chabaudi_A0A4V0K2Q1    -------------------------------I-DLNRSTVCNTEVNTKEVYLGLVCPEGY 227
P.yoelii_A0A078K5Q3      -------------------------------V-NSNKNIICNTQVNSKEVYLGLVCPEGY 228
P.vivax_A0A1G4H080       -------------------------------SLAKEKGLVCDVKITKREVYLGLVCPPGY 319
                                                            .   *:. :.  :****::** **

P.falciparum_C6KSX1      TLYPNDCFKNVIYDNNIIIPLKKI-IPHDILYHQDKNK--RITFASFTLNINENPPGFTC 321
P.berghei_Q4Z5T0         GMYPENCFENVLFEK-KVINITEL-IKHDVKLHIEKNK--NISFASFILNPNENPKSFSC 266
P.knowlesi_A0A1Y3DHK6    EMYPSNCFDRVLYKD-SIVRMSEL-IKHHVTFHMDSNR--RMSFATFSLDRNENPPGFTC 365
P.reichenowi_A0A151LP59  ILYPNDCFKNVIYDNNIIIPLKKI-IPHDILYHQDKNK--RITFASFTLNINENPPRFTC 301
P.malariae_A0A1D3SN11    TIYPSDCFENVLYEN-KIVNIKELLVSHDIKLHIDKKK--RMSFATFILNKNENPKGFSC 308
P.ovale_A0A1A8VY82       KMYPLTCFENVLHEK-EVVKINEL-VQHDVKLHMDVHK--QISFATFTLNKNENPNSFSC 276
P.chabaudi_A0A4V0K2Q1    EMYPENCFEYVLFES-NVVRINEL-IKHDVKLHIEKNKHTHMSFASFTLNPNENPKSFSC 285
P.yoelii_A0A078K5Q3      EIYPENCFENILFEN-KVIKISEL-IKHDIKLHIEKNK--NISFASFILNSNENPKSFSC 284
P.vivax_A0A1G4H080       EMYPSNCFERVLQKDGSIVRVNEL-LKHDVSFHADGNR--RMSFATFTLNRNENPQGFSC 376
                           :**  **. :: ..  :: :.:: : *.:  * : ::  .::**:* *: ****  *:*

P.falciparum_C6KSX1      YCIKDQTNINNPLIVNFHFSNQETSYATKNKNLFFYFIFIFPFLYVILLL 371
P.berghei_Q4Z5T0         HCIKNN-DNSFPLIANITFPIMNLIL-------------------LISM-- 295
P.knowlesi_A0A1Y3DHK6    LCVRMDIPEAPPLQANFVYHNYESFGFHFRL-LYVLVVIL---LLVLC-L 410
P.reichenowi_A0A151LP59  YCIKDQTNINNPLIVNFHFSNQETSYATKNTNLFFYFILIFPFLYFILFL 351
P.malariae_A0A1D3SN11    QCVKNN-DNIFPLQANFEYANYESFSLCTHL-RYFVLLLS---LLVLF-L 352
P.ovale_A0A1A8VY82       HCVKDG-DHASILQANFTYANYESASLPVRL-ALLFLLPI---LFSLL-W 320
P.chabaudi_A0A4V0K2Q1    QCIKKN-ANAFPLIANIMFSNYESYSFNYHV-TYLILISI---ILISY-I 329
P.yoelii_A0A078K5Q3      HCIKNN-DNSFPLIANITFANYESYSFNFYA-TYFILIFI---FLISY-I 328
P.vivax_A0A1G4H080       MCLNVQAPEAPPLQANFAFHNYESAGVRFGL-PCALVALV---LLALC-L 421
                          *:.         *  .*:  :    :
```

**Figure S3**

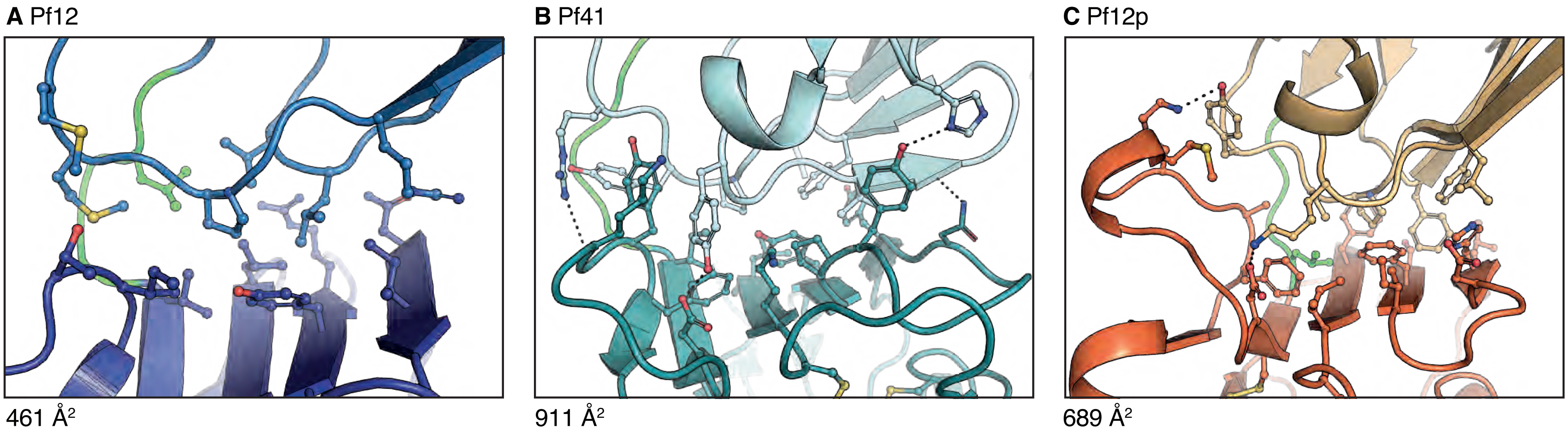
